# Heterooligomerization drives structural plasticity of eukaryotic peroxiredoxins

**DOI:** 10.1101/2025.03.03.641172

**Authors:** Jannik Zimmermann, Lukas Lang, Julia Malo Pueyo, Mareike Riedel, Khadija Wahni, Dylan Stobbe, Christopher Lux, Steven Janvier, Didier Vertommen, Svenja Lenhard, Frank Hannemann, Helena Castro, Ana Maria Tomas, Johannes M. Herrmann, Armindo Salvador, Timo Mühlhaus, Jan Riemer, Joris Messens, Marcel Deponte, Bruce Morgan

## Abstract

Peroxiredoxins are highly conserved thiol peroxidases essential for peroxide detoxification, redox signaling, and chaperone activity. Prx1/AhpC-type peroxiredoxins are found throughout the eukaryotic kingdom, where multiple isoforms frequently coexist within the same cell and even in the same subcellular compartment. Long thought to form exclusively homooligomeric structures, we reveal that heterooligomerization is a conserved and important feature of eukaryotic Prx1/AhpC-type peroxiredoxins. We demonstrate that heterooligomer formation modulates peroxoredoxin oligomeric state and enhances structural stability. In yeast, Tsa1–Tsa2 peroxiredoxin heterodecamers form in response to oxidative stress and incorporated Tsa2 stabilizes the decameric state. Beyond yeast, we show that human PRDX1 and PRDX2, as well as plant and parasitic peroxiredoxins, engage in functional heterooligomerization. These findings challenge the long-held paradigm of peroxiredoxin homooligomerization and reveal a novel mechanism for regulating redox homeostasis. Our study provides new insights into peroxiredoxin structural plasticity with broad implications for redox biology, stress responses, and cellular adaptation.

## Introduction

Peroxiredoxins are highly efficient enzymes found in nearly all living organisms, playing essential roles in maintaining cellular fitness. As some of the most abundant proteins in cells, peroxiredoxins are crucial in peroxide scavenging, redox signaling, and as molecular chaperones (**Fig. 1**) ^1–5^. These versatile enzymes are classified into various groups based on their enzymatic mechanisms or structural features.

**Figure 1.**
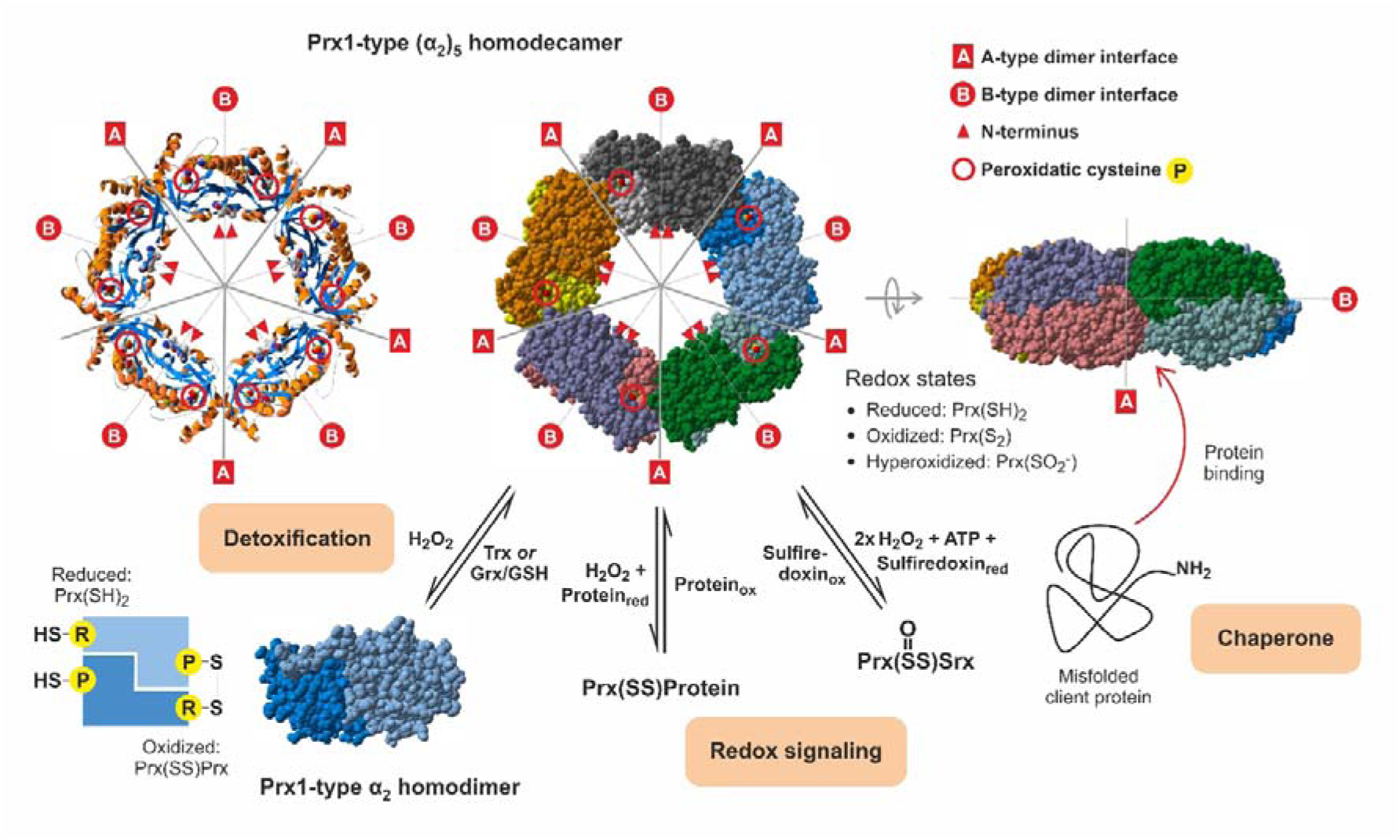
The oligomeric state of peroxiredoxin is closely linked to its function and is affected by numerous factors. Prx1/AhpC-type peroxiredoxins (Prx) exist in a dynamic equilibrium between dimeric and decameric forms form of the (α₂)₅-type depicted as a ribbon diagram (top left), space filling model (top middle), and space filling side-view (top right). The oligomeric assembly is formed through A-type (dimer– dimer) and B-type (intra-dimer) interfaces. Prx dimers (bottom left) transition between reduced [Prx(SH)₂] and oxidized [Prx(SS)] forms via reaction with H₂O₂, followed by reduction facilitated by thioredoxins (Trx) or glutaredoxins (Grx). Hyperoxidation leads to Prx(SO₂) formation, reversible via sulfiredoxin (Srx) in an ATP-dependent reaction. The function of Prx1-AhpC-type peroxiredoxins is intricately coupled to their oligomeric state and includes peroxide detoxification, redox signalling, and chaperone activity.

Peroxiredoxin-1 (Prx1)/AhpC-type peroxiredoxins are commonly found in a dynamic equilibrium between homodimers and homodecamers ^3,6,7^. Homodimers involve an interaction across the B-interface, whilst homodecamers are of the (α_2_)_5_ type, comprised of five dimers interacting via their A-type dimer interfaces (**Fig. 1**) ^8^. The dimer–decamer equilibrium is influenced by factors such as protein concentration, the redox state of the catalytic cysteine residues, pH, and various post-translational modifications ^6,9–16^. All characterized Prx1/AhpC-type peroxiredoxins are mechanistically typical 2-Cys peroxiredoxins, having two key cysteinyl residues per subunit, a peroxidatic cysteine (C_P_) and a resolving cysteine (C_R_), which are essential for catalysis ^8,17^. Dimers form through a ‘head-to-tail’ subunit arrangement, allowing for intermolecular disulfide bond formation between the C_P_ and C_R_ of the different subunits at both dimer ends (**Fig. 1**).

Many organisms harbor two or more Prx1/AhpC-type peroxiredoxins, often with two highly homologous isoforms present in the same subcellular compartment. For example, in the budding yeast *Saccharomyces cerevisiae*, Tsa1 and Tsa2 share 86% sequence identity and are both located in the cytosol ^18–20^. Likewise, human PRDX1 and PRDX2 and *Leishmania infantumLi*PRX1 and *Li*PRX2 co-exist in the cytosol and share 78% and 87% sequence identity respectively ^21–23^. Finally, in *Arabidopsis thaliana,* BAS1A and BAS1B, sharing 96% sequence identity (excluding the chloroplast-targeting transit peptides), are both located in the chloroplast stroma. Although these protein often have highly similar primary and tertiary structures, they often show significant differences in biophysical and biochemical properties including their dimer–decamer equilibrium, isoelectric points, enzyme kinetics (including C_P_ hyperoxidation kinetics), and susceptibility to various post-translational modifications ^10,24–28^.

Despite their high sequence identity, the different Prx1/AhpC-type peroxiredoxins in an organism are considered to only form homooligomeric complexes. The first challenge to the homooligomer-only model came from the characterization of peroxiredoxin-based H_2_ O_2_ sensors ^29,30^. We showed that the genetically encoded roGFP2-Tsa2ΔC_R_ probe, based on a genetic fusion between roGFP2 and the yeast peroxiredoxin Tsa2, forms enzymatically active heterooligomeric complexes with endogenous Tsa1 in yeast ^29^. More recently, human PRDX1 and PRDX2 were also shown to form heterooligomers *in vitro*, though functionality, activity, and *in vivo* relevance were not assessed ^31^. These observations led us to explore whether peroxiredoxin heterooligomerization is simply a rare phenomenon limited to a few specialized experimental settings, or if it represents a broader, biologically significant feature of eukaryotic Prx1/AhpC-type peroxiredoxins.

In this study, we reveal an exciting new aspect of peroxiredoxin biology, showing that eukaryotic peroxiredoxins can assemble into heterooligomers with a wide range of different subunit stoichiometries. Modulation of the dimer–decamer equilibrium appears to be a common outcome of heterooligomer formation between eukaryotic peroxiredoxins. In yeast, heterooligomerization between Tsa1 and Tsa2 is inducible in response to an oxidative challenge, concomitant with the induction of *TSA2* expression. The incorporation of just one Tsa2 subunit is enough to strongly stabilize a 9xTsa1–1xTsa2 decamer, consistent with a gain-of-function role for heterooligomerization driven by sub-stoichiometric amounts of one of the partner proteins. We find heterooligomerization between human PRDX1 and PRDX2 in HEK293T cells and demonstrate that *Arabidopsis* and *Leishmania* peroxiredoxins can also form functional heterooligomers. Given the relationship between peroxiredoxin oligomeric state and function, it is likely that heterooligomerization represents a prevalent and conserved mechanism for regulation and fine-tuning of peroxiredoxin function in different subcellular compartments, different cell types, and different species, throughout the eukaryotic kingdom.

## Results

### Yeast Tsa1 and Tsa2 form heterodimers and heterodecamers in Escherichia coli

We first asked whether Tsa1 and Tsa2 can form heterooligomers when purified from *E. coli* (**Fig. 2** and **Supplementary Fig. 1**). Recombinant N-terminally His_6_-or Strep-tagged Tsa1 and His_6_-tagged Tsa2 were produced and purified individually (**Supplementary Fig. 1a**) and then mixed at a 1:1 ratio. After a tandem-affinity purification using Ni-NTA agarose followed by StrepTactin agarose beads no interaction between Strep-Tsa1 and His_6_-Tsa2 was detected. This suggests that individually purified recombinant Tsa1 and Tsa2 do not exchange subunits and form stable homooligomeric complexes.

**Figure 2.**
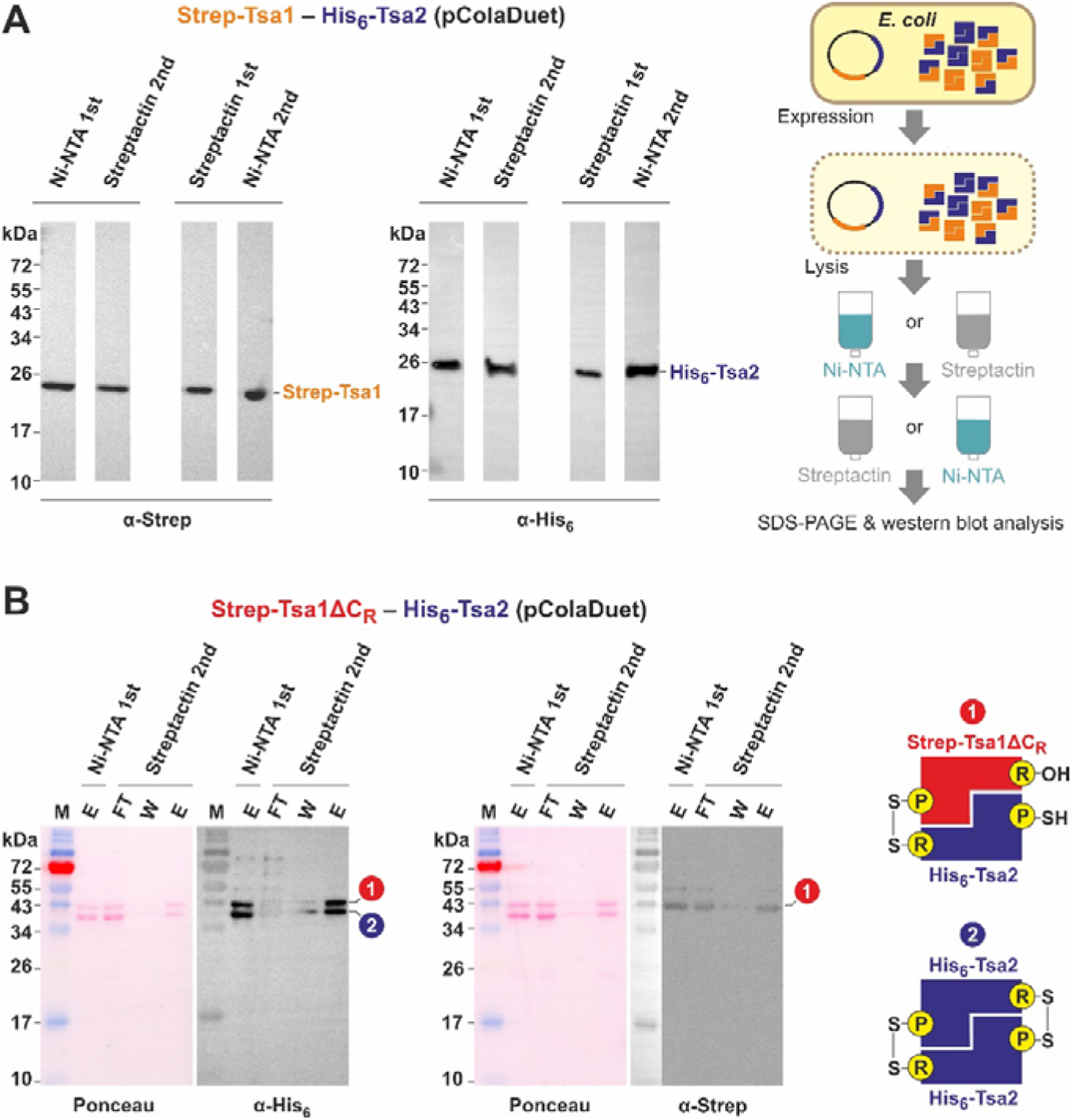
**Strep-Tsa1 and His**_6_**-Tsa2 form heterooligomers in** E. coli **A**) Purification scheme and western blot analysis of the eluates from tandem affinity co-purifications of Strep-Tsa1 and His_6_-Tsa2 with Ni-NTA agarose followed by StrepTactin agarose or *vice versa*. Eluate samples were separated by reducing SDS-PAGE. The calculated molecular masses of Strep-Tsa1 and His_6_-Tsa2 are 22.9 and 23.8 kDa, respectively. Uncropped blots are shown in Figure S1. **B**) The oligomeric state of individually or tandem-purified recombinant Tsa1 and Tsa2 was analyzed by clear-native gel electrophoresis followed by western blot analysis. **C**) Tandem affinity co-purification of Strep-Tsa1ΔC_R_ and His_6_-Tsa2. Protein samples were treated with *N*-ethylmaleimide to block free thiols and separated by non-reducing SDS-PAGE to preserve intersubunit disulfide bonds as shown in Fig. 1B. M, marker; FT, flow-through; W, wash; E, eluate

Next, we co-expressed genes encoding Strep-Tsa1 and His_6_-Tsa2 from a single plasmid in *E. coli* and performed tandem-affinity purifications with Ni-NTA agarose followed by StrepTactin agarose or *vice versa*. Under these conditions, Strep-Tsa1 and His_6_-Tsa2 were successfully co-purified regardless of the order of the tandem-affinity purification, supporting an interaction between recombinant Tsa1 and Tsa2 in *E. coli* (**Fig. 2a, Supplementary Fig. 1b,c**). Semi-quantitative western blotting, calibrated with individually purified Tsa1 and Tsa2, showed similar levels of co-purified proteins, suggesting a ∼1:1 ratio of Strep-Tsa1 and His_6_-Tsa2 (**Supplementary Fig. 1d**).

To rule out other possibilities for co-purification such as the formation of heterogenous stacks of homodecameric complexes, we performed tandem-affinity purification with the resolving cysteinyl mutant Strep-Tsa1ΔC_R_, which cannot form disulfide-linked Tsa1ΔC_R_–Tsa1ΔC_R_ homodimers, and His_6_-Tsa2, which can form either Tsa2–Tsa2 homodimers or disulfide-linked Tsa1ΔC_R_–Tsa2 heterodimers (**Fig. 2b**). Non-reducing SDS-PAGE and western blotting detected two disulfide-linked dimer species for His_6_-Tsa2, whereas a single disulfide-linked dimer species was detected for Strep-Tsa1ΔC_R_, indicating that the proteins form heterodimers through the B-type interface (**Fig. 2b**).

In conclusion, recombinant yeast Tsa1 and Tsa2 readily interact through their B-type interface, and probably also through their A-type interface, forming heterooligomeric complexes.

### Tsa1 and Tsa2 form decamers with heterogenous subunit stoichiometries

We next sought to visualize Tsa1–Tsa2 heterooligomers by negative stain electron microscopy (EM). To this end, we co-expressed Strep-Tsa1 and His_6_-Tsa2-EPEA in which Tsa2 additionally contains a C-terminal EPEA epitope, which is known to bind with high affinity to the nanobody Nbsyn2.20 ^32,33^. We produced and purified recombinant Strep-Tsa1–His_6_-Tsa2-EPEA heterooligomers using tandem Ni-NTA and StrepTactin affinity chromatography, collecting three different fractions for analysis. Strep-Tsa1 homooligomers and His_6_-Tsa2-EPEA homooligomers were purified separately as controls (**Supplementary Fig. 2a–d**). Native-PAGE analysis showed that the Tsa1–Tsa2 heterooligomers run at a mass consistent with a decamer, between the bands observed for the Tsa1 and Tsa2 homooligomers (**Supplementary Fig. 2e**) MALDI-TOF mass spectrometry (MS) analysis revealed that all Tsa1–Tsa2 heterooligomers contain disulfide-linked dimers, consistent with the lack of a reducing agent in the purification protocol (**Fig. 3a**). Notably, all three heterooligomer populations contained disulfide-linked Tsa1–Tsa2 heterodimers as well as Tsa2–Tsa2 homodimers. One of the heterooligomer fractions additionally contained disulfide-linked Tsa1–Tsa1 homodimers (**Fig. 3a**). All dimers were reducible with DTT, and no contamination of AhpC from *E. coli* was detected in the samples (**Supplementary Fig. 3a**). Further, we performed LC-MS/MS of the three purified Tsa1–Tsa2 hetero-oligomer populations revealing that each population contained different ratios of Tsa1:Tsa2 molecules: 0.25 mol/mol, 0.42 mol/mol, and 0.88 mol/mol (PRIDE accession number PXD060819) suggesting that we have heterooligomer population with different subunit stoichiometries. Finally, we analyzed Tsa1 and Tsa2 homooligomers as well as the 0.88 mol/mol Tsa1–Tsa2 heterooligomer fraction by negative stain EM showing that the Strep-Tsa1, His_6_-Tsa2-EPEA, and the Strep-Tsa1–His_6_-Tsa2-EPEA samples all contain donut-shaped structures consistent with decamers of similar diameter (**Supplementary Fig. 4**).

**Figure 3.**
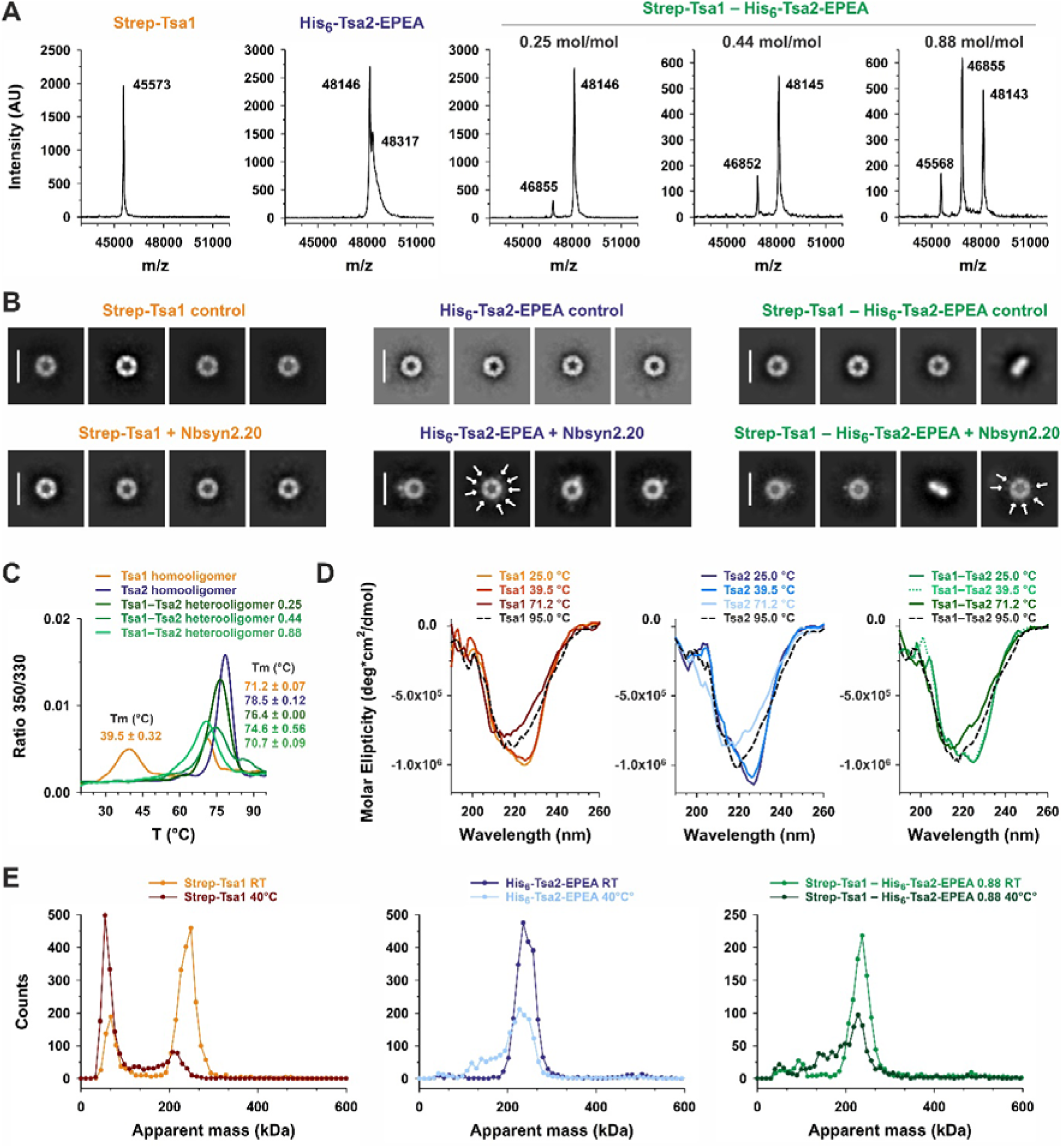
Tsa1–Tsa2 heterooligomer formation stabilizes the decameric state. **A**) MALDI-TOF mass spectrometry analysis reveals that Strep-Tsa1−His_6_-Tsa2-EPEA heterooligomers are formed by disulfide-linked homo-dimers of Strep-Tsa1 and His_6_-Tsa2-EPEA, as well as disulfide-linked heterodimers of Strep-Tsa1−His_6_-Tsa2-EPEA, with different ratios. The ratio of each Strep-Tsa1−His_6_-Tsa2-EPEA heterooligomer sample is displayed above the corresponding spectra. The signal intensity (AU) is plotted against different m/z ratios. Data were analyzed by FlexAnalysis 3.4 (Bruker). Duplicates of each sample were measured. **B**) Negative staining electron microscopy (EM) micrographs demonstrate that Nbsyn2.20 binds to 0.88 mol/mol Strep-Tsa1−His_6_-Tsa2-EPEA heterodecamer at multiple positions. Protein particles appear bright against the 2% uranyl-acetate stain, revealing a characteristic decameric’donut-like’ structure. His_6_-Tsa2-EPEA homodecamer and Strep-Tsa1−His_6_-Tsa2-EPEA heterodecamer with Nbsyn2.20 2D-classes contain one or more ‘nodules’, not present in untreated samples nor in Strep-Tsa1 homodecamer samples. Nbsyn2.20 position are marked with white arrows. Negative staining EM micrographs were analyzed using CryoSparc v4.6.0 **C**) NanoDSF experiments show that an increased presence of His_6_-Tsa2-EPEA molecules enhances the stability of Strep-Tsa1−His_6_-Tsa2-EPEA heterooligomer samples. Temperature was increased from 20°C to 100°C with a step size of 2°C/min. Strep-Tsa1 and His_6_-Tsa2-EPEA homooligomers profiles are shown in orange and blue, respectively. Strep-Tsa1−His_6_-Tsa2-EPEA heterooligomers profiles are shown in green. Triplicates of each sample were measured, and five spectra per sample were averaged. **D**) Circular dichroism (CD) data indicate no significant change in the secondary structure of Strep-Tsa1 homooligomer, His_6_-Tsa2-EPEA homooligomer, and Strep-Tsa1−His_6_-Tsa2-EPEA heterooligomersamples at 39.5°C. However, a loss of some secondary structural features occurs at higher temperatures (71.2°C for Strep-Tsa1, 78.5°C for His_6_-Tsa2-EPEA, and 70.7°C for Strep-Tsa1−His_6_-Tsa2-EPEA). The samples do not fully denature at 95°C. CD spectra of Strep-Tsa1 and His_6_-Tsa2-EPEA homooligomers are shown in orange and blue, respectively. The 0.88 mol/mol Strep-Tsa1−His_6_-Tsa2-EPEA heterooligomer CD spectra are shown in green. **E**) Mass photometry data indicate that Strep-Tsa1 homooligomer dissociates into dimers at 40°C. Mass photometry analysis of Strep-Tsa1, His_6_-Tsa2-EPEA, and the 0.88 mol/mol Strep-Tsa1–His_6_-Tsa2-EPEA sample at room temperature and 40°C were measured. Stock protein solutions (5 mM) were diluted prior to measurement. Data were acquired for 60 s, and counts of individual molecules were plotted against their molecular weight. Strep-Tsa1 and His_6_-Tsa2-EPEA homooligomers mass photometry histograms are shown in orange and blue, respectively. 0.88 mol/mol Strep-Tsa1−His_6_-Tsa2-EPEA heterooligomer mass photometry histogram is shown in green. Data were analyzed using DiscoverMP (version 2.1.1; Refeyn Ltd), with triplicates measured.

We next utilized the Nbsyn2.20 nanobody to visualize the position of Tsa2 in the heterooligomers. Biolayer Interferometry (BLI) and mass photometry confirmed the specificity of Nbsyn2.20 binding to the EPEA tag, with no binding observed to Tsa1 homooligomers (**Supplementary Fig. 5a**). Mass photometry revealed that at least two Nbsyn2.20 molecules bind to a Tsa1–Tsa2 heterodecamer. Notably, there are multiple populations of Tsa1–Tsa2 heterodecamers in solution, as indicated by an asymmetric molecular weight peak with a tail. Interestingly, the Tsa2 homooligomer control showed various populations with different numbers of Nbsyn2.20 molecules bound, likely due to the dynamic nature of the interaction (**Supplementary Fig. 5b**). Tsa1 homodecamers did not bind to Nbsyn2.20, confirming the specificity of the interaction.

Finally, negative stain EM imaging of Tsa1–Tsa2 heterodecamers in the presence of Nbsyn2.20 revealed that the nanobody binds at multiple positions on the heterodecamer, which might be explained by the existence of different stoichiometries of Tsa1 and Tsa2 in solution (**Fig. 3b**). Consistent with our BLI and mass photometry observations, we observed no interaction of Tsa1 homooligomers with Nbsyn2.20, while Tsa2 homooligomers showed up to ten Nbsyn2.20 molecules bound at various positions (**Fig. 3b**). In summary, our structural analysis of recombinant Tsa1–Tsa2 heterooligomers reveals a donut-shaped arrangement where Tsa1 and Tsa2 molecules interact in a dynamic, specific, and stoichiometry-dependent manner.

### Tsa2 stabilizes the decameric state of the heterooligomers

We next sought to evaluate the decamer stability and dimer–decamer equilibrium of Tsa1–Tsa2 heterooligomers in comparison to their homooligomeric counterparts. NanoDSF analysis of the Tsa1 homooligomer revealed two inflection points at 39.5°C and 71.2°C (**Fig. 3c**). In contrast, Tsa2 homooligomers displayed only one inflection point at 78.5°C. The three purified Tsa1–Tsa2 heterooligomer populations also presented only one inflection point at 76.4°C (0.25 mol/mol population), 74.6°C (0.42 mol/mol population), and 70.7°C (0.88 mol/mol population) (**Fig. 3c**). Our data show that the presence of more Tsa2 molecules led to increased stability of the oligomers as seen by a progressive increase in the temperature of the inflection point. Next, we performed circular dichroism (CD) measurements at temperatures of 25°C, 39.5°C, the inflection points for each sample, and 95°C (**Fig. 3d**). The CD data indicated that there is no change in the secondary structure profile at 39.5°C for any sample. In all three proteins, the second inflection point around 75°C in **Fig. 3c** indicates a loss of some secondary structural features although it is important to mention that none of the protein samples fully denature at any temperature, suggesting that the Tm values are above 95°C (**Fig. 3d**).

Finally, we used mass photometry to better understand the molecular basis of the different inflection points observed in the nanoDSF experiments (**Fig. 3e**). Mass photometry measurements were performed at room temperature and 40°C on the Tsa1 homooligomer, Tsa2 homooligomer, and 0.88 mol/mol Tsa1–Tsa2 heterooligomer samples. We found a loss of the decameric state and dissociation to dimers of Tsa1 at 40°C, supporting that the 39.5°C inflection point observed in the nanoDSF experiment reflects dissociation of Tsa1 homodecamers into homodimers. In contrast, the Tsa2 homooligomer and 0.88 mol/mol Tsa1–Tsa2 heterooligomer samples showed much less dissociation into low molecular weight oligomers (**Fig. 3e**). In summary, our data support that the Tsa1 homodecamer is less stable than the Tsa2 homodecamer, consistent with previous reports ^27^. Intriguingly, the incorporation of just a few Tsa2 subunits strongly stabilizes the decameric state of the resultant Tsa1–Tsa2 heterodecamers, suggesting that heterooligomerization might be an important mechanism for regulating peroxiredoxin oligomeric state dynamics.

### Tsa1–Tsa2 heterooligomerization is inducible and promotes decamer stabilization in yeast

Under standard growth conditions, Tsa1 is highly abundant in yeast cells, whereas Tsa2 levels are much lower ^34^. However, *TSA2* expression is known to be inducible under oxidative challenge ^35,36^. To investigate whether Tsa1 and Tsa2 heterooligomers form when expressed from their native promoters, we replaced either *TSA1* or *TSA2* in their native genomic loci with roGFP2-tagged versions, i.e. *RoGFP2-TSA1* and *RoGFP2-TSA2,* respectively, with both constructs remaining under the control of the endogenous promoter and 3’ untranslated regions. Fluorescence measurements confirmed that the roGFP2-Tsa1 level was ∼10-fold higher than that of roGFP2-Tsa2 (**Fig. 4a**). Upon exposure to 1 mM H_2_O_2_, the roGFP2-Tsa1 fluorescence remained unchanged, whereas the roGFP2-Tsa2 fluorescence increased ∼5-fold within 90 min, showing an induction of the expression of *TSA2* but not *TSA1* in response to an oxidative challenge (**Fig. 4a)**.

**Figure 4.**
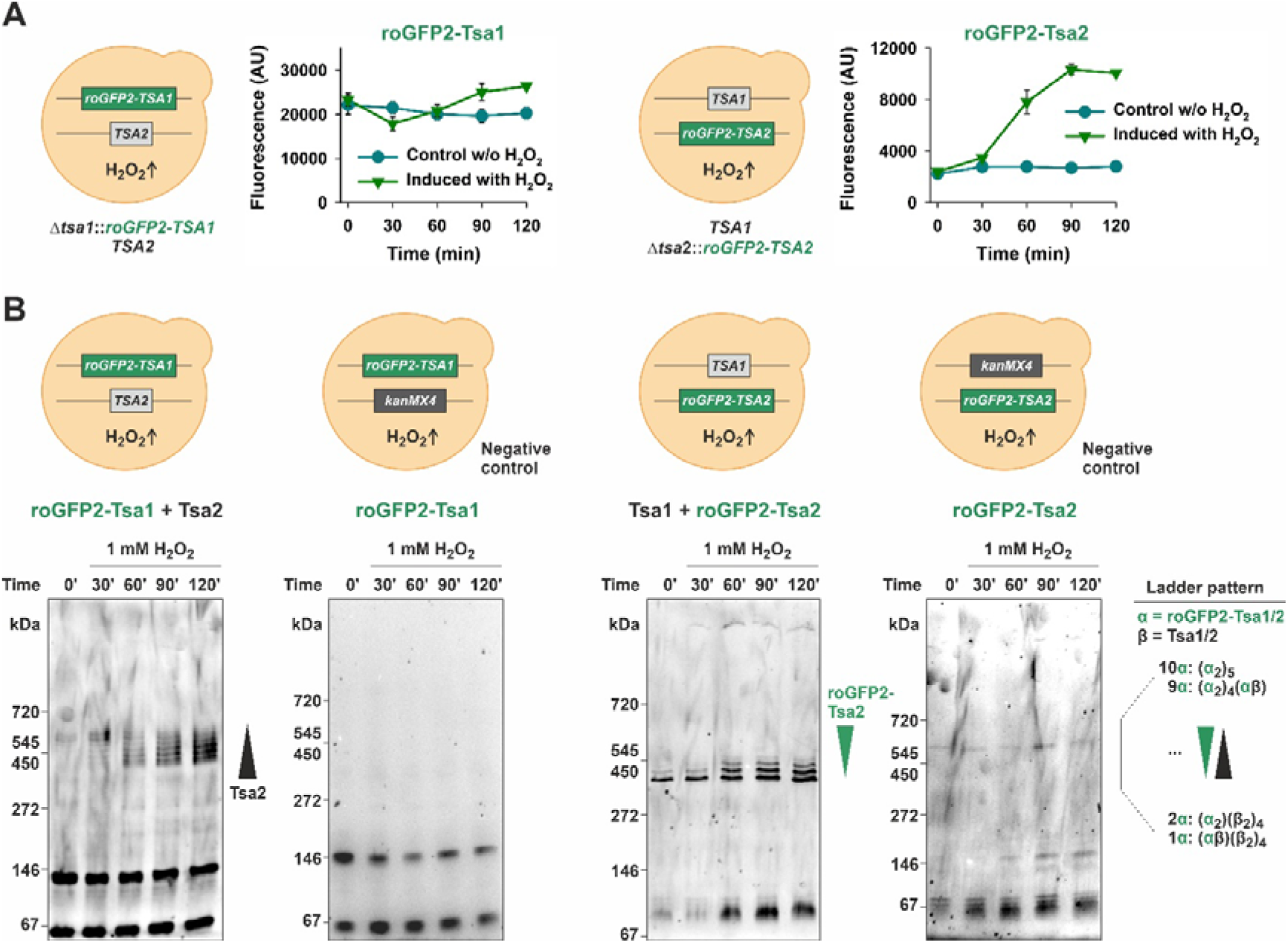
Heterooligomerization is inducible in yeast and promotes decamer stabilization A. **B**) Graph showing the GFP fluorescence intensity of **A**) roGFP2-Tsa1 and **B**) roGFP2-Tsa2 constructs in yeast cells at the indicated timepoints after treatment with 1 mM exogenous H_2_O_2_. The constructs were expressed from genes integrated into the *TSA1* and *TSA2* genomic loci respectively remaining under the control of the native promoters and 3’ untranslated regions. Experiments were repeated three times with independent yeast cultures. Data represent mean ± s.d. **C–F**) Clear-native PAGE gels, monitored for GFP fluorescence, of lysates taken from the cultures of the indicated yeast strains at the indicated timepoints following addition of 1 mM exogenous H_2_O_2_.

To further explore the impact of *TSA2* induction on heterooligomer formation, we performed Clear-Native gel electrophoresis. Under normal conditions, roGFP2-Tsa1 predominantly migrated as a dimer. Nonetheless, within 30 min after H_2_O_2_ addition to Δ*tsa1*::*RoGFP2-TSA1* cells, we observed bands at a molecular mass consistent with roGFP2-Tsa1–Tsa2 heterodecamers, which intensified during the time-course of the experiment (**Fig. 4b**). We observed a ladder pattern of decamer bands, consistent with the formation of roGFP2-Tsa1–Tsa2 heterodecamers with various subunit stoichiometries (**Fig. 4b**). No higher molecular mass bands were observed in Δ*tsa1*::*RoGFP2-TSA1* Δ*tsa2* control cells, in which the *TSA2* gene was deleted (**Fig. 4b**).

In Δ*tsa2*::*RoGFP2-TSA2* cells, the predominant species was a decamer. Upon H_2_O_2_ treatment, and subsequent induction of roGFP2-Tsa2 expression, a series of higher-molecular-mass bands appeared above the initial decamer band, suggesting that as roGFP2-Tsa2 levels increase, heterooligomers with progressively higher roGFP2-Tsa2 content form (**Fig. 4b**).

In conclusion, our findings demonstrate that Tsa1–Tsa2 heterooligomerization is inducible upon oxidative challenge and show that incorporation of just one or two Tsa2 subunits is sufficient to stabilize a heterodecameric complex.

### Heterooligomerization enables remarkable structural plasticity

Our data support the existence of Tsa1–Tsa2 interactions across both the A-and B-type interfaces, prompting us to calculate the theoretical maximum number of possible different heterooligomeric configurations. Our analysis revealed that two distinct monomer types can assemble into 120 unique decamers (i.e. structures that are not rotations of the same decamer around the 5-fold symmetry axis perpendicular to the plane of the decamer or across the five 2-fold symmetry axes parallel to this plane, see **Fig. 1a**, **Supplementary Fig. 6a** and **Supplementary Material**). These 120 distinct heterodecamers include 2 distinct homodecameric structures, 2 decamers with a 1:9 stoichiometric ratio, 14 decamers with 2:8 or 8:2 ratios, 24 decamers with 3:7 or 7:3 ratios, 52 decamers with 4:6 or 6:4 ratios, and 26 decamers with 5:5 ratio. The plasticity (defined as the ability of Prx1-type enzymes to change their structure, functions or interactions in response to their environment, including parameters such as peroxides, heat, salts, pH, and posttranslational modifications) of these heterooligomers further increases dramatically when accounting for the multiple states of the monomers. Each monomer cycles through at least three distinct structural states during the catalytic cycle, which multiply several-fold when known post-translational modifications are also considered^37^. In turn, the number of possible unique heterodecamer configurations scales with the 10^th^ power of the number of states (**Supplementary Fig. 6b and Supplementary Material**). This number would far exceed the number of peroxiredoxin decamers in a cell, assuming each of the two monomer types have only four states. The potential configurations become even more vast when more realistic numbers of states are considered. Therefore, only a minor fraction of the possible peroxiredoxin heterooligomer configurations can occur in a cell at any given moment.

### Enzymatic activity of Tsa1–Tsa2 heterooligomers mirrors that of homooligomers

We recently analyzed the catalytic cycle of recombinant His_6_-Tsa1 using stopped-flow kinetic measurements, revealing three distinct reaction phases for the H_2_O_2_-dependent oxidation of reduced Tsa1, i.e. Tsa1(SH)_2_, and three phases for the yeast Trx1-dependent reduction of Tsa1 disulfide, i.e. Tsa1(S_2_) ^38^. To compare the catalytic activity of Tsa1–Tsa2 heterooligomers to that of their homooligomeric counterparts, we analyzed individually purified His_6_-Tsa1 and His_6_-Tsa2, and co-purified mixtures of recombinant Strep-Tsa1 and His_6_-Tsa2 (**Fig. 5, Supplementary Fig. 1, Table 1).**

**Figure 5.**
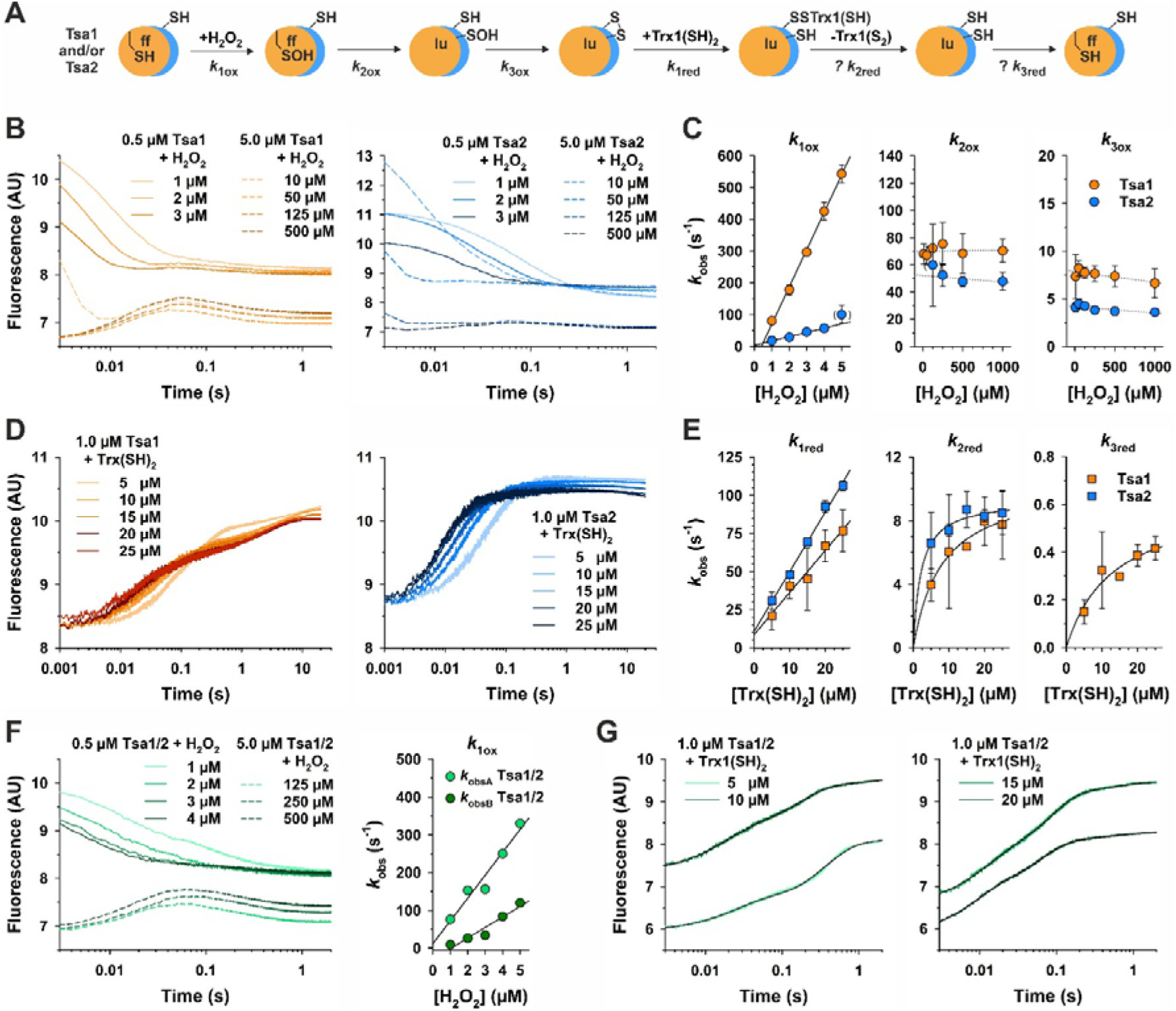
Heterooligomers of recombinant Tsa1 and Tsa2 have similar enzyme kinetics as compared to their homo-oligomers. **A**) Reaction scheme for the oxidation and reduction of recombinant Tsa1 and/or Tsa2. Only the dimeric enzyme species are shown for simplicity. **B**) Representative changes of tryptophan fluorescence during the H_2_O_2_-dependent oxidation of individually purified His_6_-Tsa1(SH)_2_ (left) or His_6_-Tsa2(SH)_2_ (right). **C**) Secondary plots of the *k*_obs_ values from exponential fits of the three reaction phases at variable H_2_O_2_ concentrations from panel **B**. **D**) Representative changes of tryptophan fluorescence during the yeast Trx1-dependent reduction of individually purified His_6_-Tsa1(S_2_) (left) or His_6_-Tsa2(S_2_) (right). **E**) Secondary plots of the *k*_obs_ values from exponential fits of the two or three reaction phases at variable Trx1 concentrations from panel **D**. **F**) Representative changes of tryptophan fluorescence during the H_2_O_2_-dependent oxidation of co-purified His_6_-Tsa1(SH)_2_ and His_6_-Tsa2(SH)_2_ (left) and secondary plot of the *k*_obs_ values from exponential fits of the first reaction phase (right). **G**) Representative changes of tryptophan fluorescence during the yeast Trx1-dependent reduction of co-purified His_6_-Tsa1(S_2_) and His_6_-Tsa2(S_2_). Quadruple exponential fits (black lines) were calculated using the *k*_obs_ values of the first phase of individually purified His_6_-Tsa1 and His_6_-Tsa2 at the according Trx1 concentration as input. All stopped-flow measurements were performed at pH 7.4 and 25°C. Ten times higher enzyme concentrations were used to highlight the second and third oxidation phase (dashed lines). Data from panels **C** and **E** were from technical triplicates of independent biological triplicate protein purifications and measurements. Data from panel **F** and fits in panel **G** were from technical triplicates of a single measurement. Rate constants are summarized in Table 1.

**Table 1.** Rate constants for the redox reactions of recombinant Tsa1 and/or Tsa2 at pH7.4 and 25°C.

| Protein | Assigned reaction partners A/B and products P/Q |  |  |  | Rate constant |  |
| --- | --- | --- | --- | --- | --- | --- |
|  | A | + B | P | + Q | k |  |
| Tsa1 | His<sub>6</sub>-Tsa1(SH)<sub>2</sub> | H<sub>2</sub>O<sub>2</sub> | His<sub>6</sub>-Tsa1(SOH)<sub>ff</sub> | H<sub>2</sub>O | k<sub>1ox</sub> | (1.2 ± 0.03) × 10<sup>8</sup> M<sup>-1</sup>s<sup>-1</sup> |
| Tsa1 | His<sub>6</sub>-Tsa1(SH)<sub>2</sub><sup>a</sup> | H<sub>2</sub>O<sub>2</sub> | His<sub>6</sub>-Tsa1(SOH)<sub>ff</sub><sup>a</sup> | H<sub>2</sub>O | k<sub>1ox</sub> | (9.7 ± 0.4) × 10<sup>7</sup> M<sup>-1</sup>s<sup>-1</sup> <sup>a</sup> |
| Tsa1 | Strep-Tsa1(SH)<sub>2</sub> | H<sub>2</sub>O<sub>2</sub> | Strep-Tsa1(SOH)<sub>ff</sub> | H<sub>2</sub>O | k<sub>1ox</sub> | 8.1 × 10<sup>7</sup> M<sup>-1</sup>s<sup>-1</sup> <sup>b</sup> |
| Tsa2 | His<sub>6</sub>-Tsa2(SH)<sub>2</sub> | H<sub>2</sub>O<sub>2</sub> | His<sub>6</sub>-Tsa2(SOH)<sub>ff</sub> | H<sub>2</sub>O | k<sub>1ox</sub> | (1.3 ± 0.1) × 10<sup>7</sup> M<sup>-1</sup>s<sup>-1</sup> |
| Tsa1/2 | His<sub>6</sub>-Tsa1(SH)<sub>2</sub>/ | H<sub>2</sub>O<sub>2</sub> | His<sub>6</sub>-Tsa1(SOH)<sub>ff</sub>/ | H<sub>2</sub>O | k<sub>1oxA</sub> | 6.1 × 10<sup>7</sup> M<sup>-1</sup>s<sup>-1</sup> |
|  | Strep-Tsa2(SH)<sub>2</sub> |  | Strep-Tsa2(SOH)<sub>ff</sub> |  | k<sub>1oxB</sub> | 2.8 × 10<sup>7</sup> M<sup>-1</sup>s<sup>-1</sup> |
| Tsa1+Tsa2 | His<sub>6</sub>-Tsa1(SH)<sub>2</sub> + | H<sub>2</sub>O<sub>2</sub> | His<sub>6</sub>-Tsa1(SOH)<sub>ff</sub> + | H<sub>2</sub>O | k<sub>1oxA</sub> | 9.8 × 10<sup>7</sup> M<sup>-1</sup>s<sup>-1</sup> |
|  | His<sub>6</sub>-Tsa2(SH)<sub>2</sub> |  | His<sub>6</sub>-Tsa2(SOH)<sub>ff</sub> |  | k<sub>1oxB</sub> | 1.2 × 10<sup>7</sup> M<sup>-1</sup>s<sup>-1</sup> |
| Tsa1 | His<sub>6</sub>-Tsa1(SOH)<sub>ff</sub> |  | His<sub>6</sub>-Tsa1(SOH)<sub>lu</sub> |  | k<sub>2ox</sub> | 70 ± 9 s<sup>-1</sup> |
| Tsa1 | His<sub>6</sub>-Tsa1(SOH)<sub>ff</sub><sup>a</sup> |  | His<sub>6</sub>-Tsa1(SOH)<sub>lu</sub><sup>a</sup> |  | k<sub>2ox</sub> | 64 ± 5 s<sup>-1</sup> <sup>a</sup> |
| Tsa1 | Strep-Tsa1(SOH)<sub>ff</sub> |  | Strep-Tsa1(SOH)<sub>lu</sub> |  | k<sub>2ox</sub> | 67 s<sup>-1</sup> <sup>b</sup> |
| Tsa2 | His<sub>6</sub>-Tsa2(SOH)<sub>ff</sub> |  | His<sub>6</sub>-Tsa2(SOH)<sub>lu</sub> |  | k<sub>2ox</sub> | 51 ± 11 s<sup>-1</sup> |
| Tsa1/2 | His<sub>6</sub>-Tsa1(SOH)<sub>ff</sub>/ |  | His<sub>6</sub>-Tsa1(SOH)<sub>lu</sub>/ |  | k<sub>2ox</sub> | 45 ± 7 s<sup>-1</sup> |
|  | Strep-Tsa2(SOH)<sub>ff</sub> |  | Strep-Tsa2(SOH)<sub>lu</sub> |  |  |  |
| Tsa1 | His<sub>6</sub>-Tsa1(SOH)<sub>lu</sub> |  | His<sub>6</sub>-Tsa1(S<sub>2</sub>) | H<sub>2</sub>O | k<sub>3ox</sub> | 6.6 ± 1.7 s<sup>-1</sup> |
| Tsa1 | His<sub>6</sub>-Tsa1(SOH)<sub>lu</sub><sup>a</sup> |  | His<sub>6</sub>-Tsa1(S<sub>2</sub>)<sup>a</sup> | H<sub>2</sub>O | k<sub>3ox</sub> | 4.6 ± 0.3 s<sup>-1</sup> <sup>a</sup> |
| Tsa1 | Strep-Tsa1(SOH)<sub>lu</sub> |  | Strep-Tsa1(S<sub>2</sub>) | H<sub>2</sub>O | k<sub>3ox</sub> | 6.8 s<sup>-1</sup> <sup>b</sup> |
| Tsa2 | His<sub>6</sub>-Tsa2(SOH)<sub>lu</sub> |  | His<sub>6</sub>-Tsa2(S<sub>2</sub>) | H<sub>2</sub>O | k<sub>3ox</sub> | 4.0 ± 0.1 s<sup>-1</sup> |
| Tsa1/2 | His<sub>6</sub>-Tsa1(SOH)<sub>lu</sub>/ |  | His<sub>6</sub>-Tsa1(S<sub>2</sub>)/ | H<sub>2</sub>O | k | 3.6 ± 0.6 s<sup>-1</sup> |
|  | Strep-Tsa2(SOH)<sub>lu</sub> |  | Strep-Tsa1(S<sub>2</sub>) |  |  |  |
| Tsa1 | His<sub>6</sub>-Tsa1(S<sub>2</sub>)<sup>b</sup> | Trx1(SH)<sub>2</sub> | His<sub>6</sub>-Tsa1-SS-Trx1<sup>b</sup> |  | k<sub>1red</sub> | (2.8 ± 0.5) × 10<sup>6</sup> M<sup>-1</sup>s<sup>-1</sup> |
| Tsa2 | His<sub>6</sub>-Tsa2(S<sub>2</sub>) | Trx1(SH)<sub>2</sub> | His<sub>6</sub>-Tsa2-SS-Trx1 |  | k<sub>1red</sub> | (3.9 ± 0.1) × 10<sup>6</sup> M<sup>-1</sup>s<sup>-1</sup> |
| Tsa1 | ? His<sub>6</sub>-Tsa1-SS-Trx1 |  | ? His<sub>6</sub>-Tsa1(SH)<sub>2</sub> lu | Trx1(S<sub>2</sub>) | k<sub>2red</sub> | 10 ± 2 s<sup>-1</sup> |
| Tsa2 | ? His<sub>6</sub>-Tsa2-SS-Trx1 |  | ? His<sub>6</sub>-Tsa2(SH)<sub>2</sub> lu | Trx1(S<sub>2</sub>) | k<sub>2red</sub> | 9.4 ± 0.7 s<sup>-1</sup> |
| Tsa1 | ? His<sub>6</sub>-Tsa1(SH)<sub>2</sub> lu |  | ? His<sub>6</sub>-Tsa1(SH)<sub>2</sub> ff |  | k<sub>3red</sub> | 0.6 ± 0.1 s<sup>-1</sup> |

<sup>a</sup> 39, <sup>b</sup> 38

As previously reported for Tsa1 ^27,38,39^, three distinct phases were observed during the oxidation of individually purified His_6_-Tsa1(SH)_2_ and His_6_-Tsa2(SH)_2_: (i) a decrease of tryptophan fluorescence reflecting the rapid formation of the sulfenic acid species at higher H_2_O_2_ concentrations, (ii) an increase in fluorescence indicating a conformational change from the fully folded (ff) to the locally unfolded (lu) state, and (iii) a subsequent decrease in fluorescence due to the formation of an intermolecular disulfide bond at the B-type dimer interface (**Fig. 5a,b**). While the second-order rate constant *k*_1ox_ of 1.3 × 10^7^ M^-1^s^-1^ for the sulfenic acid formation of His-Tsa2 was ten times smaller than for His_6_-Tsa1, the first-order rate constants (*k*_2ox_ and *k*_3ox_) for the H_2_O_2_-independent phases were similar for both enzymes (**Fig. 5c**, **Table 1**). Control experiments with Strep-Tsa1 revealed that the Strep-tag had no major effect on the oxidation kinetics compared to His_6_-Tsa1 (**Table 1**).

For the yeast Trx1-dependent reduction, three phases were observed for His_6_-Tsa1(S_2_) and two phases for His-Tsa2(S_2_) (**Fig. 5a,d**). The second-order rate constant *k*_1red_ of 3.9 × 10^6^ M^-1^s^-1^ for the formation of the mixed disulfide between His_6_-Tsa2 and yeast Trx1 was slightly higher than that for the mixed disulfide of His-Tsa1 (**Fig. 5e**, **Table 1**). The similar y-axis intercepts at 8.7 s^-1^ for His-Tsa1 and at 10.8 s^-1^ for His_6_-Tsa2 suggest a reverse reaction or a conformational change. The second and third phases revealed pseudo-first-order reaction kinetics at higher Trx1 concentrations with similar rate constants *k*_2red_ ∼10 s^-1^ for both His-Tsa1 and His-Tsa2 (**Fig. 5e**, **Table 1**). The second phase likely corresponds to the formation of reduced peroxiredoxins and Trx1(S_2_). The third phase, with <u>k</u>_3red_ (0.6 s^-1^) for His-Tsa1 may indicate local refolding, a process not observed for His-Tsa2.

The oxidation kinetics of co-purified recombinant Strep-Tsa1 and His_6_-Tsa2 were generally similar to those of the individual enzymes (**Fig. 5f**) but required a quadruple exponential fit to characterize the first oxidation phase. The rate constants, *k* and *k*, had intermediate values of 6.1 × 10^7^ M^-1^s^-1^ and 2.8 × 10^7^ M^-1^s^-1^ compared to the *k*_1ox_ values of the individual enzymes (**Fig. 5f, Tab l**)**e**. T**1**he second and third phase displayed similar kinetics to those observed for individual enzymes, yielding *k* and *k* values of 45 and 3.6 s^-1^, respectively. Pre-mixed mixtures of His-Tsa1 and His-Tsa2 at a 1:1 ratio showed similar oxidation kinetics as the individual enzymes, with *k*_1oxA_ and *k*_1oxB_ values from a double exponential fit of 9.8 × 10^7^ M^-1^s^-1^ and 1.2 × 10^7^ M^-1^s^-1^ (**Table 1**). Thus, the oxidation kinetics of Strep-Tsa1 and His_6_-Tsa2 mixtures reflect the superposition of the individual enzyme activities, whereas co-purified heterooligomers of Strep-Tsa1 and His_6_-Tsa2 have altered, intermediate macroscopic rate constants (*k*_1ox_) for the reaction with H_2_O_2_.

The reduction kinetics of co-purified recombinant Strep-Tsa1 and His_6_-Tsa2 resembled those of His_6_-Tsa1 but required a quadruple exponential fit with two pre-set rate constants (**Fig. 5g**). Best results were obtained using *k*_obs_ values for the first reduction phase from the individually purified enzymes at the corresponding Trx1 concentration. The remaining *k*_obs_ values were consistent with those observed for His_6_-Tsa1. This suggests that the reduction kinetics of the co-purified heterooligomers of Strep-Tsa1 and His_6_-Tsa2 can be approximated by superimposing the reduction profiles of the individually purified enzymes.

In summary, the oxidation and reduction kinetics of individually purified, mixed, and co-purified recombinant Tsa1 and Tsa2 revealed similar enzyme kinetics for heterooligomers compared to their homooligomeric counterparts. Although the formation of the sulfenic acid species in His_6_-Tsa1 was ten times faster than in His_6_-Tsa2, co-purified Tsa1–Tsa2 heterooligomers exhibited intermediate macroscopic rate constants for the reaction with H_2_O_2_.

### Tsa1 and Tsa2 form enzymatically active heterooligomers in yeast

We previously demonstrated that a roGFP2-Tsa2ΔC_R_ probe can form enzymatically active heterooligomers with endogenous Tsa1 in the yeast cytosol ^29^. To further investigate the assembly and enzymatic activity of Tsa1–Tsa2 heterooligomers in yeast, we expressed roGFP2-Tsa1 variants in Δ*tsa1*Δ*tsa2* yeast cells alongside either wild-type (wt) or catalytically inactive Tsa2. Specifically, cells were transformed with plasmids encoding roGFP2-Tsa1ΔC_P_ΔC_R_, in which both catalytic cysteine residues were mutated to serine, and either Tsa2wt or Tsa2ΔC_P_ΔC_R_. Cells were then exposed to increasing concentrations of H_2_O_2_ (0–500 µM), and roGFP2 oxidation was quantified using a fluorescence plate-reader assay (**Supplementary Fig. 7**).

In Δ*tsa1*Δ*tsa2* cells co-expressing roGFP2-Tsa1ΔC_P_ΔC_R_ and Tsa2wt, roGFP2 oxidation was detected at exogeneous H_2_O_2_ concentrations as low as 10 µM, indicating a highly sensitive response. In contrast, no roGFP2 oxidation was observed in cells co-expressing roGFP2-Tsa1ΔC_P_ΔC_R_ and Tsa2ΔC_P_ΔC_R_, even at the highest H_2_O_2_ concentration. Since roGFP2 is only very poorly directly oxidized by H₂O_2_, these results confirm that Tsa1 and Tsa2 assemble into functional heterooligomers in the yeast cytosol ^29,30,40,41^, where roGFP2 oxidation is catalyzed *in trans* by a neighboring peroxiredoxin within the complex. A similar experiment using a roGFP2-Tsa2ΔC_P_ΔC_R_ co-expressed with either Tsa1wt or Tsa1ΔC_P_ΔC_R_ yielded comparable results (**Supplementary Fig. 7**). Efficient roGFP2 oxidation was observed in the presence of Tsa1wt and almost no oxidation in the presence of Tsa1ΔC_P_ΔC_R_.

In summary, our findings confirm that Tsa1 and Tsa2 can form catalytically active heterooligomers in the yeast cytosol, functioning cooperatively to mediate roGFP2 oxidation.

### Native human PRDX1 and PRDX2 form heterooligomers in HEK293 cells

To determine whether peroxiredoxin heterooligomerization extends beyond yeast, we investigated human PRDX1 and PRDX2, which were recently shown to form heterooligomers *in vitro* ^31^. Despite their 78% sequence identity, PRDX1 and PRDX2 have distinct isoelectric points (7.80 and 5.71, respectively), allowing us to utilize ion-exchange chromatography to separate them in Flp-In™ T-Rex™ HEK293 cell lysates (**Fig. 6**). Western blot analysis of collected fractions revealed that although PRDX1 and PRDX2 primarily eluted at different points along the NaCl gradient, they also eluted across a broad range of overlapping fractions (**Fig. 6a,b**).

**Figure 6.**
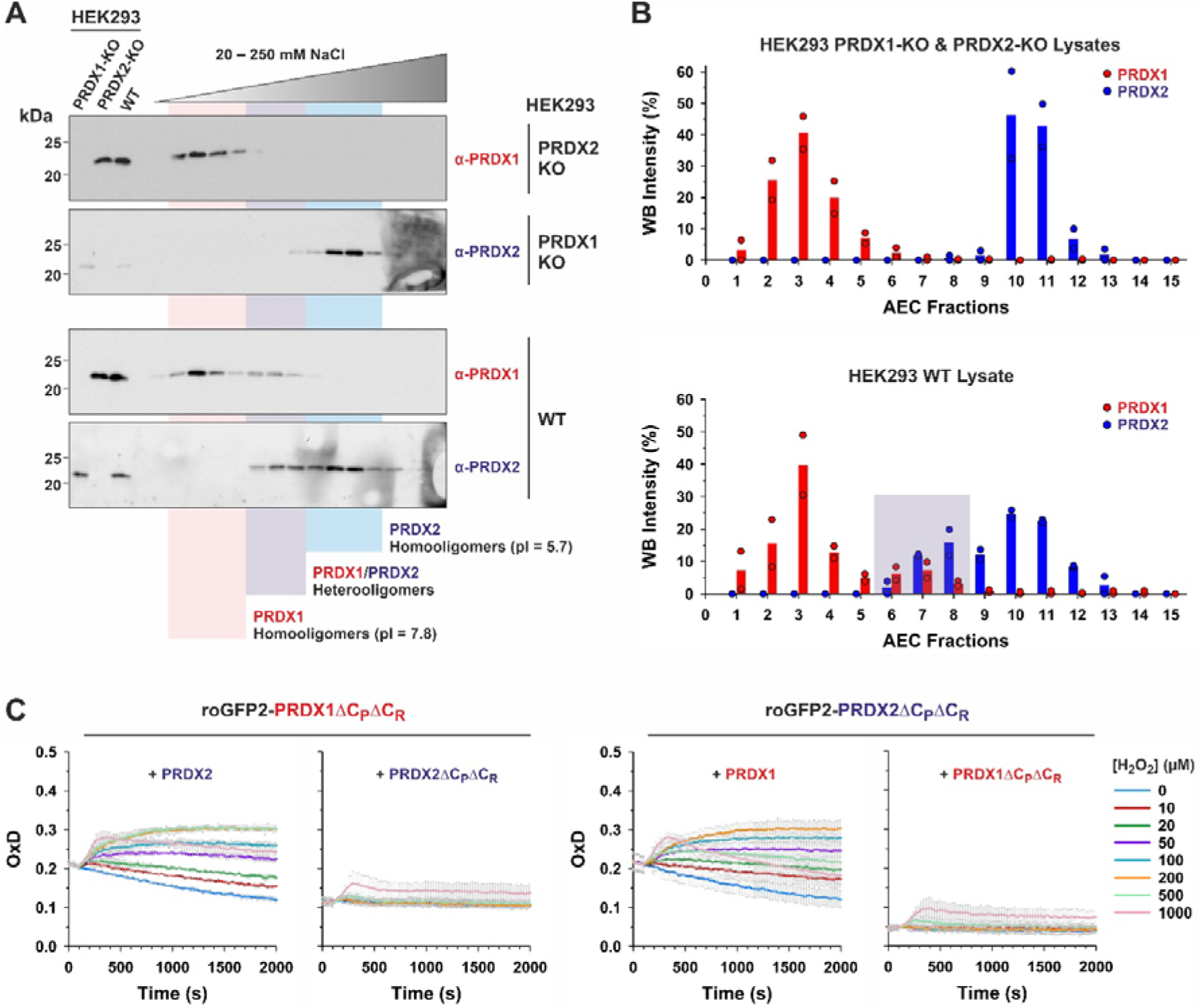
Native human PRDX1 and PRDX2 form heterooligomers In HEK293 cells. **A**) Western blot analysis and **B**) quantification of anion exchange chromatography fractions from HEK293 WT and PRDX1 and PRDX2 KO cell lysates. Proteins from the three cell lines were separated by anion exchange chromatography and fractions subsequently analyzed by Western blot for the presence of PRDX1 and PRDX2. Lysates from the three cell-lines were prepared and fractionated independently, with no mixing of lysates at any stage. Western blots (left panel) and quantification (right panel) show that in the PRDX KO cell lines the PRDX 1 and 2 signals show no overlap in the middle fractions, whereas in the WT lysate there is a clear overlap in the middle fractions, indicating a shift in the isoelectric point of the protein complex species, suggesting the presence of heterooligomers. **C**) Graphs showing change in degree of oxidation (OxD) of the indicated roGFP2-PRDX fusion constructs in Δ*tsa1*Δ*tsa2* yeast cells treated with the indicate concentrations of H_2_O_2_. Experiments were repeated 3 times with independent yeast cultures. Data are presented as mean ± s.d.

To further explore this interaction, we repeated the experiment using lysates from CRISPR-Cas9 knockout (KO) cells lacking either PRDX1 or PRDX2 ^42^. In PRDX1-KO and PRDX2-KO cells, the remaining PRDX isoform eluted in a narrower range of fractions, with no overlap observed (**Fig. 6a,b**). This shift in elution pattern suggests that PRDX1 and PRDX2 influence each other’s chromatographic behavior, providing strong evidence for their interaction as native heterooligomers.

To assess whether PRDX1–PRDX2 heterooligomers are enzymatically active, we employed a yeast-based roGFP2 assay. We produced heterologous roGFP2-PRDX1ΔC_P_ΔC_R_ in Δ*tsa1*Δ*tsa2* yeast cells alongside either PRDX2 or PRDX2ΔC_P_ΔC_R_ and measured roGFP2 oxidation upon H₂O₂ exposure (**Fig. 6c**). Consistent with our observation with yeast Tsa1 and Tsa2, roGFP2 oxidation was observed only when PRDX2 was present, indicating that PRDX1 requires an active PRDX2 partner for mediating roGFP oxidation. Similarly, roGFP2-PRDX2ΔC_P_ΔC_R_ only supported roGFP2 oxidation when co-expressed with PRDX1, but not PRDX1ΔC_P_ΔC_R_ (**Fig. 6c**).

In summary, our findings demonstrate that PRDX1 and PRDX2 form enzymatically active heterooligomers in both HEK293T cells and yeast and support the presence of peroxiredoxin heterooligomerization across species.

### Peroxiredoxins from diverse eukaryotes form enzymatically active heterooligomers

To further assess whether heterooligomerization is a conserved feature of peroxiredoxins in other eukaryotes, we examined peroxiredoxins from non-opisthokont organisms using the yeast-based roGFP2 system. We generated roGFP2 fusions constructs with *Li*PRX1 and *Li*PRX2 from the cytosol of the kinetoplastid parasite *Leishmania infantum* ^23^, as well as BAS1A and BAS1B from the chloroplast stroma of the plant *Arabidopsis thaliana*^43,44^. Each construct was expressed in Δ*tsa1*Δ*tsa2* yeast cells together with either the wild-type of double cysteine mutant of the corresponding peroxiredoxin partner (**Fig. 7a**). In all cases, roGFP2 oxidation was observed exclusively in the presence of the wild-type partner, but not with the cysteine mutant, consistent with our observations with yeast and human peroxiredoxins.

**Figure 7.**
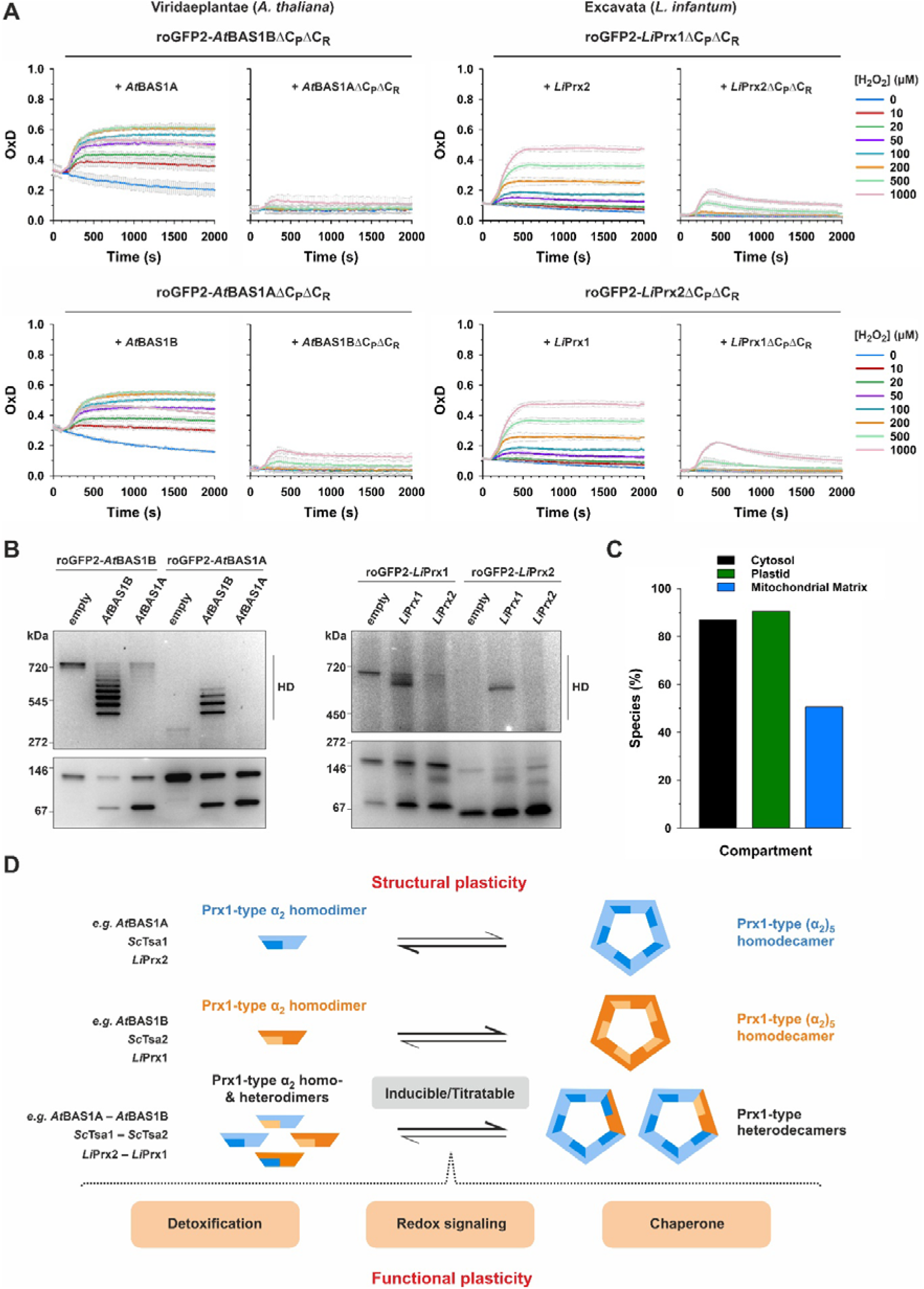
Heterooligomerization is a common feature of eukaryotic peroxiredoxins. **A**) Graphs showing the change in the degree of oxidation (OxD) in response to 1 mM H_2_O_2_ of roGFP2-Tsa1ΔC_P_ΔC_R_ and roGFP2-Tsa1ΔC_P_ΔC_R_ constructs expressed in Δ*tsa1*Δ*tsa2* yeast together with either a wild-type (wt) or cysteine-less (ΔC_P_ΔC_R_) variant of the corresponding partner peroxiredoxin. Experiments were repeated 3 times with independent yeast cultures. Data are presented as mean ± s.d. **B**) Clear-native PAGE analysis of lysates from Δ*tsa1*Δ*tsa2* yeast strains expressing the indicated combination of constructs. Gels were imaged for GFP fluorescence. Gels presented are representative examples of three independent experimental replicates. **C**) Graph showing the percentage of sequenced eukaryotic genomes in which there are predicted to be two or more Prx1/AhpC-type peroxiredoxin present in the indicated subcellular compartments. **D**) Model illustrating the impact of peroxiredoxin heterooligomerization on oligomeric state. Substoichiomeitry subunit incorporation is capable of strongly shifting the dimer–decamer equilibrium.

To explore the oligomeric state of these heterooligomers, we co-expressed *roGFP2-BAS1A* and *roGFP2-BAS1B* with either an empty vector, *BAS1A*, or *BAS1B* (**Fig. 7b**). While roGFP2-BAS1B predominantly formed a decamer, roGFP2-BAS1A was almost exclusively dimeric. Co-expression of *BAS1A* with *roGFP2-BAS1B*induced a shift from a decamer to a heterodimeric form, whereas co-expression of *BAS1B* with *roGFP2-BAS1B* maintained decamer stability, producing a series of heterodecamers with varying stoichiometries and a minor heterodimer band. Conversely, co-expression of *BAS1B* with *roGFP2-BAS1A* led to partial decamer formation, but only when BAS1A subunits were a minority, supporting that BAS1A destabilizes decamers. Co-expression of *roGFP2-BAS1A* and *BAS1A* only led to the formation of heterodimers.

Similar results were observed with *L. infantum* peroxiredoxins (**Fig. 7b**). In yeast lysates, roGFP2-LiPrx1 primarily formed decamers in contrast to roGFP2-LiPrx2, which did not oligomerize. Co-expression of *LiPRX2* with *roGFP2-LiPRX1* disrupted decamer formation, promoting a shift toward putative heterodimeric species, whereas co-expression with *LiPRX1* did not affect decamer stability. Consistent with the apparent difference in decamer stabilities, co-expression of *roGFP2-LiPRX2* with *LiPRX1* induced a shift to decameric species, whilst co-expression with *LiPRX2* did not.

In summary, our experiments suggest that strong differences in dimer–decamer equilibria are widespread among eukaryotic peroxiredoxins. Even minimal incorporation of a second peroxiredoxin subunit can dramatically alter the dominant oligomeric state, highlighting the functional significance of heterooligomerization across diverse organisms.

### Pairs of peroxiredoxins are present in most sequenced eukaryotic genomes

Finally, we sought to determine the percentage of eukaryotes in which two or more Prx1/AhpC-type peroxiredoxin proteoforms are predicted to be present in the same subcellular compartment. Extensive sequence search resulted in 11,415 peroxiredoxin candidate sequences covering 1,525 eukaryotic species across the kingdoms Viridaeplantae, Metazoa and Fungi. Of these species, 1,471 were predicted to contain at least one peroxiredoxin in the cytosol, with two or more cytosolic peroxiredoxins predicted in 1,326 species (**Fig. 7c and Supplementary Fig. 8**). Mitochondria-targeted peroxiredoxins were predicted in 1,236 species with 771 species predicted to contain two or more mitochondrial peroxiredoxins. Within the kingdom Viridaeplantae, plastid-localized peroxiredoxins were predicted in 347 species, with two or more plastid-localized peroxiredoxin in 314 species.

Overall, our data indicate that peroxiredoxin heterooligomerization is likely to be relevant in more than 90% of sequenced eukaryotic species and probably occurs in several different subcellular compartments.

## Discussion

We reveal that heterooligomerization is a common feature of Prx1/AhpC-type peroxiredoxins throughout the eukaryotic kingdom. Peroxiredoxins have been intensively studied since their discovery more than 30 years ago with thousands of publications on these fascinating proteins ^1,3,5,7,45,46^. Nonetheless, until now peroxiredoxins have been almost exclusively considered to form homooligomeric complexes. Our findings necessitate a careful reevaluation of peroxiredoxin structural dynamics and open new avenues for exploring their functional diversity (**Fig. 7d**).

The biological function of peroxiredoxin heterooligomerization remains an open question. While a complete answer remains elusive, our data strongly suggest that modulation of the dimer–decamer equilibrium plays a critical role. Previous research indicates that different oligomeric states are preferentially associated with different cellular activites. For example, the decameric state of Prx1/AhpC-type peroxiredoxins is associated with chaperone activity ^47–51^, while dimers are associated with peroxidase function ^6,11,52^. Higher-order assemblies, including stacked decamers and 12-decamer-containing dodecahedrons, may contribute to additional cellular functions yet to be fully understood ^13,53^.

Interestingly, differences in dimer–decamer equilibria seem to be a common feature among different peroxiredoxins within an organism. For example, human PRDX1 is more stable as a decamer than PRDX2 ^10,24^. Similarly, yeast Tsa2 is more stable in the decameric form than Tsa1 ^27^. Our data further support a strong difference in dimer–decamer equilibria of *A. thaliana* chloroplast and *L. infantum* cytosolic peroxiredoxins. However, we now must strongly consider the possibility that this binary classification of peroxiredoxins as either ‘type-A’ or ‘type-B’ based on their oligomeric preferences is probably an oversimplification. This view, largely derived from studies of recombinant proteins *in vitro*, may not accurately reflect the dynamic environment inside living cells. Instead, in the absence of mechanisms to actively prevent heterooligomerization — of which we have found no evidence — cells are likely to contain a complex mixture of peroxiredoxin heterooligomers with diverse subunit stoichiometries, each exhibiting a different dimer–decamer equilibrium (**Fig. 7d**).

The incorporation of just one Tsa2 subunit appears to be sufficient to stabilize a 9xTsa1–1xTsa2 decamer, with the progressive incorporation of more Tsa2 subunits leading to further stabilization. The structural basis for this effect remains unclear. However, in the crystal structure of Tsa2 decamer, the C-termini of most subunits are not visible in the electron density ^54^. This contrasts strongly with the Tsa1 decamer crystal structures in which all C-termini are clearly resolved ^55^. Furthermore, in our reduction kinetics analysis, we did not observe a third phase — attributed to local refolding — for Tsa2, unlike Tsa1. We speculate that these structural and kinetic differences may be important for the significant enhanced decamer stability of Tsa2 homodecamers and Tsa1– Tsa2 heterodecamers compared to Tsa1 homodecamers.

Prx1/AhpC-type peroxiredoxins within the same organism can also vary in several physico-chemical parameters including isoelectric point, enzyme kinetics, sensitivity to hyperoxidation, and susceptibility to various post-translational modifications including limited proteolysis ^56^, tyrosine nitration ^57^, peroxidatic cysteine *S*-nitrosylation ^58^, *S*-gluathionylation ^14^, and phosphorylation ^59^. As a result, beyond simply modulating the dimer–decamer equilibrium, heterooligomerization can also generate peroxiredoxin oligomers with a wide range of different properties. These variations likely influence not only enzyme kinetics but also likely modulate interaction with target proteins in redox signaling relays and chaperone client proteins. In summary, the structural plasticity of peroxiredoxins could reflect their functional plasticity, and peroxiredoxins might serve as a general integration hub for compartment-specific signal transduction pathways (**Fig. 7d**).

Peroxiredoxins are important transducers and transmitters in redox signaling, facilitating the oxidation of specific target proteins. For example, PRDX2 was reported to transmit oxidation to the transcription factor STAT3 ^60,61^, whilst PRDX1 oxidizes various proteins, including the MAP kinase ASK1 ^62^. However, this raises an important question: How can a specific peroxiredoxin isoform selectively oxidize a target protein if most cells likely do not contain exclusively peroxiredoxin homooligomers? In other words, what are the implications of heterooligomerization for peroxiredoxin-dependent redox signaling? Understanding how mixed oligomers influence target specificity and signalling dynamics will be essential for unravelling the full complexity of peroxiredoxin function.

Are heterooligomers exclusive to Prx1-type enzymes? In principle, Prx1-type enzymes share a high degree of structural similarity with Prx6-type enzymes in contrast to other peroxiredoxin subfamilies^8^. To explore this possibility, we recently analyzed the cytosolic Prx1-and Prx6-type enzymes from the malaria parasite *Plasmodium falciparum* but found no evidence of interaction ^63^. In summary, while further studies are needed to investigate potential interactions among different peroxiredoxin subfamilies, current evidence suggests that the formation of heterodimers and heterooligomers is likely restricted to members of the Prx1/AhpC-type subfamily.

Future studies should explore how heterooligomerization affects peroxiredoxin-mediated signaling pathways and whether it plays a role in disease contexts where redox homeostasis is disrupted. Given the growing recognition of peroxiredoxins in aging, cancer, and neurodegenerative diseases, understanding how heterooligomerization influences their function could provide new insights into redox regulation and potential therapeutic targets. Furthermore, it will be important to determine the structural determinants of peroxiredoxin heterooligomerization. Cryo-EM and crystallographic approaches could provide high-resolution insights into the interaction interfaces that stabilize heterooligomers stability, while targeted mutagenesis studies could identify key residues that drive heterooligomer formation and determine their functional consequences.

In summary, peroxiredoxin heterooligomerization represents a largely underexplored layer of regulatory mechanism with profound implications for both cellular function and organismal health.

## Materials and Methods

### Materials

N,N,N′,N′-Tetramethylazodicarboxamide (diamide), dithiothreitol (DTT), ethylenediaminetetraacetic acid (EDTA), and *N*-ethylmaleimide (NEM) were purchased from Sigma-Aldrich, diethylenetriaminepentaacetic acid (DTPA) was from Carl Roth, isopropyl-*β*-D-1-thiogalactopyranoside was from Serva, H_2_O_2_ was from VWR, and desthiobiotin was from IBA Lifesciences. PCR primers were purchased from Metabion. All restriction enzymes, DNA polymerase, and T4 DNA ligase were from New England Biolabs (NEB).

### Cloning and site-directed mutagenesis

All primers and constructs are listed in **Supplementary Table 1**. *TSA1* was PCR-amplified with Phusion HF DNA polymerase using pET15b/His-*TSA1* as a template and subcloned into the *Kpn*I and *Avr*II restriction sites of pET45b/Strep-*PFPRX1a*^C50S^ yielding pET45b/Strep-*TSA1*. Strep-*TSA1* was subsequently PCR-amplified and subcloned into the *Nde*I and *Xho*I restriction sites of the MCS2 of pColaDuet (Novagen) yielding pColaDuet/Strep-*TSA1*. Plasmid pET15b/His-*TSA2* was digested with *Nco*I and *Bam*HI and subcloned into the according restriction sites of the MCS1 of pColaDuet/Strep-*TSA1* to obtain pColaDuet/His-*TSA2*/Strep-*TSA1*. The resolving cysteine mutant of Tsa1 *TSA1ΔC*_R_ was generated by site-directed mutagenesis with Phusion HF DNA polymerase using pET45b/Strep-*TSA1* as a template yielding pET45b/Strep-*TSA1ΔC*_R_. Strep-*TSA1ΔC*_R_ was subcloned into pColaDuet together with His-*TSA2* as described above yielding pColaDuet/His-*TSA2*/Strep-*TSA1ΔC*_R_. The EPEA tag-encoding sequence was introduced into *TSA2* by PCR using pColaDuet/His-*TSA2*/Strep-*TSA1* as a template and subcloned into the *Nco*I and *Bam*HI restiction sites of the same plasmid to obtain pColaDuet/His-*TSA2*-EPEA/Strep-*TSA1*. His-*TSA2*-EPEA was subcloned into the *Nco*I and *Bam*HI restriction sites of pET15b/His-*TSA2* yielding pET15b/His-*TSA2*-EPEA. All PCR products were transformed into chemically competent *Escherichia coli* XL1-Blue cells and plasmids were isolated by minipreparation. Correct sequences were confirmed for all inserts by Sanger sequencing (Microsynth Seqlab).

### Heterologous expression and protein purification (Figure 2)

All proteins are listed in **Supplementary Table 1**. Recombinant N-terminally MGSSH_6_SSGLVPRGSHM-tagged Tsa1 and Tsa2, N-terminally MAWSHPQFEKGGT-tagged Tsa1 and pColaDuet-encoded constructs were produced in *E. coli* strain SHuffle T7 express (NEB) at 30°C. Recombinant N-terminally MH_6_P-tagged ScTrx1 was produced in *E. coli* strain XL1-Blue at 37°C. Expression of the constructs was induced with 0.5 mM IPTG for 4 h. Cultures were subsequently cooled in an ice-bath for 15 min and centrifuged (3750×*g*, 15 min, 4°C). Cell pellets were resuspended in ice-cold buffer I (20 mM imidazole, 100 mM Na_x_H_y_PO_4_, 300 mM NaCl, pH 8.0 at 4°C) and stirred on ice for 45 min with DNaseI and 10 mg lysozyme (per 1 L *E. coli* culture) before sonication and centrifugation (10000×*g*, 30 min, 4°C). Supernatants containing His-tagged Tsa1, Tsa2, or ScTrx1 from 1 L *E. coli* culture were loaded on 0.5 mL equilibrated Ni-NTA agarose columns (Qiagen). The columns were washed with 15 column volumes (CV) of ice-cold buffer I before proteins were eluted with ice-cold buffer II (200 mM imidazole, 100 mM Na_x_H_y_PO_4_, 300 mM NaCl, pH 8.0 at 4°C). Supernatants containing Strep-tagged Tsa1 from 1 L *E. coli* culture were loaded on 1 mL equilibrated StrepTactin Superflow columns (IBA Lifesciences). The columns were washed with 5 CV of ice-cold Strep-buffer I (100 mM Tris/HCl, 150 mM NaCl, 1 mM EDTA, pH 8.0 at 4°C) before proteins were eluted with ice-cold Strep-buffer II (100 mM Tris/HCl, 150 mM NaCl, 1 mM EDTA, 1 mM desthiobiotin, pH 8.0 at 4°C). For the tandem purification using pColaDuet constructs, supernatants were first purified by Ni-NTA affinity chromatography and different fractions were then seperately purified by StrepTactin affinity chromatography as described above. For the reverse tandem purification, supernatants were first purified by StrepTactin affinity chromatography and dfferent fractions were then seperately purified by Ni-NTA affinity chromatography. The puritiy of all proteins was confirmed by analytical SDS-PAGE and protein concentrations were determined spectrophotometrically at 280 nm.

### Tsa1–Tsa2-EPEA heterooligomer expression and purification (Figure 3)

pCOLA-Duet-1 plasmid, containing Strep-Tsa1 and His_6_-Tsa2-EPEA, was expressed in T7-Shuffle *E. coli* strain. 1 L LB media with kanamycin (50 μg/mL) was inoculated with a 100-fold dilution of an overnight pre-culture and grown at 30°C with shaking at 160 rpm until the exponential growth phase was reached (OD_600_ = 0.4-0.6). Then, 0.5 mM IPTG was added, and the culture was grown over night at 30°C with shaking at 160 rpm.

Cells were pelleted by centrifugation at 3750 x g for 15 min at 4°C and resuspended in lysis buffer (100 mM HEPES/NaOH pH 7.9, 300 mM NaCl, 1 μg/mL DNaseI, 50 μg/mL leupeptin, and 0.1 mg/mL AEBSF). Cells were lysed by sonication (70% amplitude for 10 min, 10” ON, 10” OFF) at 4°C, and centrifuged at 39,846 x g for 25 min at 4°C. Supernatant was filtered through a 0.45 μm filter and incubated with previously equilibrated Ni^2+^-Sepharose High Performance (Cytiva) beads for 1 h at 4°C in equilibration buffer (100 mM HEPES/NaOH pH 7.9, and 300 mM NaCl). Proteins were eluted using a stepwise gradient of 100 mM HEPES/NaOH pH 7.9, 300 mM NaCl and 1 M imidazole. Fractions containing StrepII-Tsa1 and His-Tsa2-EPEA were dialyzed overnight at 4°C against 100 mM Tris/HCl pH 8, 150 mM NaCl, and 1 mM EDTA with 2 buffer changes.

The dialyzed sample was loaded onto a Streptactin column (IBA Lifesciences) equilibrated with 100 mM Tris/HCl pH 8, 150 mM NaCl, and 1 mM EDTA, then eluted with the same buffer plus 1 mM desthiobiotin in a 1-step gradient. Fractions containing the heterooligomer consisting of StrepII-Tsa1 and His-Tsa2-EPEA were dialyzed against 100 mM Tris/HCl pH 8, 150 mM NaCl, and 1 mM EDTA. Protein concentration was determined spectroscopically using an extinction coefficient of 29,575 M^−1^ cm^−1^ for StrepII-Tsa1 homo-oligomer, 24,075 M^−1^ cm^−1^ for His-Tsa2-EPEA homo-oligomer and 26,825 M^−1^ cm^−1^ for StrepII-Tsa1/His-Tsa2-EPEA hetero-oligomer. Proteins were stored at −80°C.

Proteins were validated by Western blot. SDS-PAGE (4-20% gradient) (Biorad) separated samples were transferred to polyvinyl difluoride (PVDF) membranes (Thermo Fisher) using the wet/tank blotting system (Bio-Rad Laboratories). Membranes were blocked with 5% milk Tris-buffered saline with 0.1% Tween20 (TBS-T) for 2 h at room temperature (RT), then probed with anti-EPEA (Proteogenix), HRP-conjugated anti-StrepII (Sigma) and anti-His (Biorad) primary antibodies in 5% milk in TBS-T and incubated overnight at 4°C. Membranes were washed three times with TBS-T for 5 min shaking at RT. Then, anti-EPEA and anti-His membranes were incubated with HRP-conjugated goat anti-mouse IgG secondary antibody (Thermo Fisher) in 3% milk in TBS-T for 1 h at RT with shaking. Membranes were washed three times with TBS-T for 5 min shaking at RT. Protein bands were visualized with HRP substrate Pierce™ ECL Western Blotting Substrate (Thermo Fisher).

### Tsa1 homooligomer and Tsa2-EPEA homooligomer expression and purification (Figure 3)

pET45b plasmid, containing Strep-Tsa1, and pET15b, containing His_6_-Tsa2-EPEA, were independently expressed in T7-Shuffle *E. coli* strain. 1 L LB media with ampicillin (100 μg/mL) was inoculated with a 100-fold dilution of an overnight pre-culture and grown at 30°C with shaking at 160 rpm until the exponential growth phase was reached (OD_600_ = 0.4-0.6). Then, 0.5 mM IPTG was added, and the culture was grown over night at 30°C with shaking at 160 rpm.

Cells were pelleted by centrifugation at 3750 x g for 15 min at 4°C and resuspended in lysis buffer (100 mM HEPES/NaOH pH 7.9, 300 mM NaCl, 1 μg/mL DNaseI, 50 μg/mL leupeptin, and 0.1 mg/mL AEBSF). Cells were lysed by sonication (70% amplitude for 10 min, 10” ON, 10” OFF) at 4°C, and centrifuged at 39,846 x g for 25 min at 4°C. Supernatant was filtered through a 0.45 μm filter.

For His_6_-Tsa2-EPEA, supernatant was incubated with previously equilibrated Ni^2+^-Sepharose High Performance (Cytiva) beads for 1 h at 4°C in equilibration buffer (100 mM HEPES/NaOH pH 7.9, and 300 mM NaCl). Proteins were eluted using a stepwise gradient of 100 mM HEPES/NaOH pH 7.9, 300 mM NaCl and 1 M imidazole. Fractions containing His-Tsa2-EPEA were dialyzed against 100 mM Tris/HCl pH 8, 150 mM NaCl, and 1 mM EDTA. Protein concentration was determined spectroscopically using an extinction coefficient of 24,075 M^−1^ cm^−1^ for His-Tsa2-EPEA homo-oligomer. Proteins were stored at −80°C.

For Strep-Tsa1, supernatant was loaded onto a StrepTactin column (IBA Lifesciences) equilibrated with 100 mM Tris/HCl pH 8, 150 mM NaCl, and 1 mM EDTA, then eluted with the same buffer plus 1 mM desthiobiotin in a 1-step gradient. Fractions containing the heterooligomer consisting of Strep-Tsa1 were dialyzed against 100 mM Tris/HCl pH 8, 150 mM NaCl, and 1 mM EDTA. Protein concentration was determined spectroscopically using an extinction coefficient of 29,575 M^−1^ cm^−1^ for Strep-Tsa1 homo-oligomer. Proteins were stored at −80°C.

Proteins were validated by Western blot. SDS-PAGE (4-20% gradient) (Biorad) separated samples were transferred to polyvinyl difluoride (PVDF) membranes (Thermo Fisher) using the wet/tank blotting system (Bio-Rad Laboratories). Membranes were blocked with 5% milk Tris-buffered saline with 0.1% Tween20 (TBS-T) for 2 h at room temperature (RT), then probed with anti-EPEA (Proteogenix), HRP-conjugated anti-StrepII (Sigma) and anti-His (Biorad) primary antibodies in 5% milk in TBS-T and incubated overnight at 4°C. Membranes were washed three times with TBS-T for 5 min shaking at RT. Then, anti-EPEA and anti-His membranes were incubated with HRP-conjugated goat anti-mouse IgG secondary antibody (Thermo Fisher) in 3% milk in TBS-T for 1 h at RT with shaking. Membranes were washed three times with TBS-T for 5 min shaking at RT. Protein bands were visualized with HRP substrate Pierce™ ECL Western Blotting Substrate (Thermo Fisher).

### Sample preparation and stopped-flow measurements

Freshly purified proteins were reduced with 5 mM DTT for 30 min on ice. Excess imidazole and DTT were removed on a PD-10 desalting column (Merck). The reduced proteins were eluted in ice-cold assay buffer (100 mM Na_x_H_y_PO_4_, 0.1 mM DTPA, pH 7.4 at 25°C) and protein concentrations determined spectrophotometrically at 280 nm. Oxidized Tsa1 and Tsa2 were generated by incubation of the reduced enzyme with equimolar H_2_O_2_ for 30 min on ice. Stopped-flow measurements were performed in a thermostatted SX-20 spectrofluorometer (Applied Photophysics) at 25 °C (total emission at an excitation wavelength of 295 nm, slit width 2 mm). The oxidation of the peroxiredoxins was measured by mixing 1 µM or 10 µM of reduced Tsa1 and/or Tsa2 in syringe 1 with different concentrations of H_2_O_2_ in syringe 2. The reduction of oxidized peroxiredoxins by ScTrx1 was measured by mixing 2 µM oxidized Tsa1 and/or Tsa2 in syringe 1 with different concentrations of reduced ScTrx1 in syringe 2. The traces of at least 3 technical replicates were averaged and fitted by double or triple exponential regression using the Pro-Data SX software (Applied Photophysics) to obtain *k*_obs_ values for each phase. The *k*_obs_ values of 3 biological replicates were plotted against the substrate concentrations in SigmaPlot 13.0 to obtain second order rate constants from the slope of the linear fits or first order rate constants from linear or hyperbolic fits.

### Western blot analysis

Purified protein samples were heated for 5 min at 95°C in Laemmli buffer and seperated via SDS-PAGE or clear native PAGE (Serva) under reducing or non-reducing conditions. Samples containing tandem-purified His-TSA2 and Strep-TSA1*ΔC*_R_ were incubated with 10 mM NEM for 1 h on ice before the addition of Laemmli buffer to prevent the formation of artifical disulfide bonds. The proteins were blotted onto a PVDF membrane and stained with ponceau. Destained membranes were probed with a monoclonal mouse anti-His antibody (Invitrogen, MA1-21315) or mouse anti-Strep antibody (IBA Lifesciences, 2-1507-001) and a commercially available goat anti-mouse antibody (BioRad). The concentration of Tsa1 and Tsa2 in the tandem purification was calculated from the greyscales of the western blot bands in ImageJ as compared to different concentrations of isolated Strep-Tsa1 and His-Tsa2 that were determined spectrophotometrically.

### Nbsyn2.20 expression and purification

His-Nbsyn2.20 was expressed and purified as described ^32^, with the addition of a size exclusion step on a Superdex75 10/300 (Cytiva) equilibrated in PBS pH 7.4 as the final step. Fractions containing Nbsyn2.20 were collected and protein concentration was determined spectroscopically using an extinction coefficient of 27,180 M^−1^ cm^−1^. Nbsyn2.20 was stored at-20°C.

His-Nbsyn2.20 was validated by Western blot. The protein sample was separated by SDS-PAGE (4-20% gradient) (Biorad), transferred to polyvinyl difluoride (PVDF) membranes (Thermo Fisher) using the wet/tank blotting system (Bio-Rad Laboratories). Membranes were blocked with 5% milk Tris-buffered saline with 0.1% Tween20 (TBS-T) shaking for 2 h at RT, then probed with anti-His antibody (Biorad) primary antibody in 5% milk in TBS-T with shaking overnight at 4°C. Membranes were washed three times with TBS-T for 5 min shaking at RT. After three 5-min washes with TBS-T at RT, membranes were incubated with HRP-conjugated goat anti-mouse IgG secondary antibody (Thermo Fisher) in 3% milk in TBS-T for 1 h at RT with shaking. Membranes were washed three times with TBS-T for 5 min shaking at RT. Protein bands were visualized by the HRP substrate Pierce™ ECL Western Blotting Substrate (Thermo Fisher).

### Bilayer Interferometry (BLI)

For the BLI assay on Octet® R8 system (Sartorius), Strep-Tsa1 homooligomer, His_6_-Tsa2-EPEA homooligomer, Strep-Tsa1−His_6_-Tsa2-EPEA heterooligomer, CA17998-EPEA nanobody (positive control), and BSA (negative control) were biotinylated. Proteins were biotinylated using EZ-Link™ NHS-LC-Biotin (Thermo Scientific) following the manufacturer’s instructions. Prior to biotinylation, proteins were dialyzed into PBS pH 7.4. A 10 mM stock of NHS-LC-Biotin was prepared as a 10 mM stock solution in DMSO was mixed with protein samples at 10:1 molar ratio (NHS-LC-Biotin:protein) and incubated for 2 h on ice. Excess NHS-LC-biotin was removed by dialysis in 10 mM HEPES/NaOH pH 8, 137 mM NaCl, 2 mM KCl using Slide-A-Lyzer cassettes (10K MWCO) (Thermo Fisher) with two buffer changes, followed by overnight dialysis at 4°C. Protein concentration was determined spectroscopically using an extinction coefficient of 29,575 M^−1^ cm^−1^ for Strep-Tsa1 homooligomer, 24,075 M^−1^ cm^−1^ for His-Tsa2-EPEA homooligomer, 26,825 M^−1^ cm^−1^ for the Strep-Tsa1−His-Tsa2-EPEA heterooligomer, 25,565 M^−1^ cm^−1^ for protein-EPEA, and 43,824 M^−1^ cm^−1^ for BSA. All proteins were prepared in 10 mM HEPES/NaOH pH 8, 137 mM NaCl, and 2 mM KCl, in a final concentration of 10 μg/mL, and loaded onto the Streptavidin (SA) Biosensors (Sartorius) for 100 s at 25°C. The concentration of Nbsyn2.20 (analyte) was fixed at 50 nM in 10 mM HEPES/NaOH pH 8, 137 mM NaCl, 2 mM KCl, 1% BSA and 0.05% Tween20. Association and dissociation phases were recorded in real-time at 25°C (600 s each step). Data analysis was conducted using Sartorius Octet Analysis Studio 13.0 software. Baseline adjustments, reference subtraction, and Savitzky–Golay filtering were applied, followed by local fitting (model 1:1) to determine association and dissociation kinetic parameters. Triplicates of each sample were measured.

### Nano-Differential Scanning Fluorimetry (nanoDSF) Prometheus

Strep-Tsa1 homooligomer, His_6_-Tsa2-EPEA homooligomer, and Strep-Tsa1−His_6_-Tsa2-EPEA heterooligomer were dialyzed for 1 h against 10 mM HEPES/NaOH pH 8, 137 mM NaCl, 2 mM KCl using Slide-A-Lyzer™ Dialysis Cassettes (10K MWCO) (Thermo Fisher) with 2 buffer changes at 4°C. Protein concentrations were determined spectroscopically using an extinction coefficient of 29,575 M^−1^ cm^−1^ (Strep-Tsa1 homooligomer), 24,075 M^−1^ cm^−1^ (His_6_-Tsa2-EPEA homooligomer), 26,825 M^−1^ cm^−1^ (Strep-Tsa1−His_6_-Tsa2-EPEA heterooligomer). Protein samples (5 μM) were prepared in dialysis buffer and stored at room temperature. Inflection points were determined by monitoring fluorescence emission at 330 nm and 350 nm during a heat ramp (100°C, rate 2°C/min), with the fluorescence intensity ratio (350/330 nm) used to calculate melting temperatures from the inflection point of the denaturation curves. Triplicates of each sample were measured.

### Circular Dichroism (CD)

Strep-Tsa1 homooligomer, His_6_-Tsa2-EPEA homooligomer and Strep-Tsa1−His_6_-Tsa2-EPEA heterooligomer were initially buffer-exchanged in 10 mM sodium phosphate pH 8, 140 mM NaF buffer using a Hitrap® desalting column (Cytiva). The concentration of the proteins was determined spectroscopically using an extinction coefficient of 1.291 L g^−1^ cm^−1^ for StrepII-Tsa1 homooligomer, 0.995 L g^−1^ cm^−1^ for His-Tsa2-EPEA homooligomer and 1.143 L g^−1^ cm^−1^ for Strep-Tsa1−His-Tsa2-EPEA heterooligomer. Proteins were diluted in phosphate buffer to reach a protein concentration of 0.4 mg/mL for Tsa1 and Tsa2 homooligomers, and 0.35 mg/mL for Tsa1−Tsa2 heterooligomers. Circular dichroism measurements were performed using a BioLogic MOS-500 circular dichroism spectropolarimeter (BioLogic, France) at different temperatures (based on inflection points on nanoDSF measurements) in a quartz cuvette with a 1-mm path length (Hellma Analytics). Far UV spectra (190 to 260 nm) were recorded at a scan speed of 50 nm/min, 1 nm bandwith. Five spectra were measured and averaged. CD signal for Strep-Tsa1 homooligomer was recorded at 25°C, 39.5°C, 71.2°C and 95°C. CD signal for His-Tsa2-EPEA homooligomer was recorded at 25°C, 39.5°C, 78.5°C and 95°C. CD signal for Strep-Tsa1−His_6_-Tsa2-EPEA heterooligomer was recorded at 25°C, 39.5°C, 70.7°C and 95°C.

### Mass photometry

5 mM Strep-Tsa1 homooligomer, His_6_-Tsa2-EPEA homooligomer and Strep-Tsa1−His_6_-Tsa2-EPEA hetero-oligomer samples were incubated at room temperature and 40°C for 15 min. Protein landing was recorded using a Refeyn OneMP (Refeyn Ltd) MP system by adding 10 μL of a 25x dilution of the protein stock solution (5 mM) into a 10 μL drop of filtered buffer 10 mM HEPES/NaOH pH 8, 137 mM NaCl, and 2 mM KCl. Movies (6,000 frames, 60 s) were acquired with the AcquireMP (version 2.1.1; Refeyn Ltd) software using default settings. Data were analysed using default settings on DiscoverMP (version 2.1.1; Refeyn Ltd). Contrast-to-mass calibration was performed with MassFerence P1 (Refeyn) using standards of 88, 172, 258, and 344 kDa. The binding and unbinding events were grouped into mass ranges (binning) using GraphPad prism. Frequency distribution with default setting and a bin width of log_2_(x), being x the total number of detected particles, was used. Data was represented as number of particles (counts) vs. mass (kDa). Triplicates of each sample were measured, and the average molecular weight and standard deviation were calculated.

### Tsa1−Tsa2 heterooligomer−Nbsyn2.20 complex reconstruction

Strep-Tsa1−His_6_-Tsa2-EPEA heterooligomer and Nbsyn2.20 samples were stored at-80°C in 100 mM Tris/HCl pH 8, 150 mM NaCl, and-20°C in 1x PBS pH 7.4, respectively. The Strep-Tsa1−His_6_-Tsa2-EPEA heterooligomer was dialyzed for 1 h against 10 mM HEPES/NaOH pH 8, 137 mM NaCl, 2 mM KCl using a Slide-A-Lyzer™ Dialysis Cassettes (10K MWCO) (ThermoFisher) with 2 buffer changes at 4°C. Protein concentration was determined spectroscopically using an extinction coefficient of 26,825 M^−1^ cm^−1^ for the Strep-Tsa1−His-Tsa2-EPEA heterooligomer and 27,180 M^−1^ cm^−1^ for Nbsyn2.20. A 5 μM solution of Strep-Tsa1−His_6_-Tsa2-EPEA heterooligomer was incubated with 6 μM of Nbsyn2.20 for 20 min at RT prior to analysis by mass photometry and negative-staining electron microscopy (EM).

### Negative staining electron microscopy (EM)

Formvar/Carbon 400 Mesh, Cu grids (Electron Microscopy Sciences) were glow discharged at 4-5 mA and 0.3 mbar vacuum for 30 s. 3 μL of freshly diluted sample (0.02 mg/mL) in a buffer containing 10 mM HEPES/NaOH pH 8, 137 mM NaCl, 2 mM KCl was incubated on the grids for 30 s, followed by staining with 2% uranyl-acetate.

20 micrographs were collected on a JEOL 1400+ microscope, equipped with a LaB6 filament operating at 120 kV. Micrograph were recorded using a TVIPS F416 CCD camera using a nominal magnification of 60,000, corresponding to a magnified pixel size of 1.94 Å/px and a defocus range of 0 to-1.5 μm. The micrographs were processed using CryoSparc v4.6.0., After running patch CTF estimation, particles were picked by blob-picker and extracted using a box size of 256 px. These particles were subjected to 2D classification. Further, particle diameters were measured using EMMenu from TVIPS imaging software.

### Matrix-assisted laser desorption ionization–time of flight (MALDI-TOF) mass spectrometry (MS)

Matrix Assisted Laser Desorption Ionization – Time of Flight MS data was acquired on a Ultraflextreme enhanced MALDI TOF/TOF-MS system (Bruker Daltonics, Bremen, Germany) using FlexControl 3.4 acquisition software (Bruker).

All homooligomer and heterooligomer samples were buffer exchanged with 0.1% TFA prior to MALDI-TOF analysis. To obtain monomers, protein samples were mixed with dithiothreitol (DTT) in a 1 to 20 molar ratio, incubated for 15 min at room temperature and subsequently buffer exchanged to a 0.1% v/v TFA solution with 1 mM DTT added. Subsequently, 1 µL of a 1:1:1 mixture of protein sample, 2.5 DHAP matrix (Bruker) and 2% v/v TFA was spotted in triplicate on an MTP ground steel plate. After crystallization, spectra were measured in the linear positive ion-mode within a mass range of 5 000 to 50 000 m/z. Up to 12 000 shots were acquired with a laser repetition rate of 2000 Hz and 200 shots per rasterspot. All acquisition methods were provided by the manufacturer and optimized and calibrated with in-house made calibration standard (22 to 44 kDa, 4 calibrants). The obtained spectra were analyzed and processed (peak picking, smoothing and baseline substraction) with FlexAnalysis 3.4 (Bruker). Duplicates of each sample were measured.

### LC-MS/MS of purified Tsa1/Tsa2 oligomers

Purified proteins (40 µg) were precipitated with Methanol/Chloroform then resuspended in 40 µl of 50 mM NH_4_HCO_3_ and digested overnight with trypsin at 37°C, peptides were quantified with Pierce Quantitative Colorimetric assay (ref 23275) Peptides were dissolved in solvent A (0.1% trifluoroacetic acid in 2% acetonitrile) containing 100 ftmol/µl of digested CytC (Thermo Scientific ref 161089),100ng peptides were directly loaded onto reversed-phase pre-column (Acclaim PepMap 100, Thermo Scientific) and eluted in backflush mode. Peptide separation was performed using a reversed-phase analytical column (PepMap RSLC, 0.075 x 250 mm EasySpray,Thermo Scientific ES902) with a linear gradient of 4%-27.5% solvent B (0.1% formic acid in 80% acetonitrile) for 40 min, 27.5%-50% solvent B for 20 min, 50%-95% solvent B for 10 min and holding at 95% for the last 10 min at a constant flow rate of 300 nL/min on an Vanquish Neo system. The peptides were analyzed by an Orbitrap Fusion Lumos mass spectrometer (Thermo Fisher Scientific). The peptides were subjected to NSI source followed by tandem mass spectrometry (MS/MS) in Orbitrap Lumos coupled online to nano-LC. Intact peptides were detected in the Orbitrap at a resolution of 120,000. Peptides were selected for MS/MS using HCD setting at 30, ion fragments were detected in the IonTrap. A data-dependent procedure that alternated between one MS scan followed by MS/MS scans was applied for 3 seconds for ions above a threshold ion count of 1.0×10^4^ in the MS survey scan with 30.0s dynamic exclusion. MS1 spectra were obtained with an automatic gain control target of 4×10^5^ ions and a maximum injection time set to auto, and MS2 spectra were acquired with an automatic gain control target of 1×10^4^ ions and a maximum injection set to auto. For MS scans, the *m/z* scan range was 350 to 1800. The resulting MS/MS data was processed using Sequest HT search engine within Proteome Discoverer 2.5 SP1 against an *Escherichia coli* K12 protein database obtained from Uniprot, the sequences of Tsa1 and Tsa2 recombinant proteins and that of pig CytC. Trypsin was specified as a cleavage enzyme allowing up to 2 missed cleavages, 4 modifications per peptide and up to 5 charges. Mass error was set to 10 ppm for precursor ions and 0.2 Da for-fragment ions. Oxidation on Met (+15.995 Da), conversion of Gln (-17.027 Da) or Glu (-18.011 Da) to pyro-Glu at the peptide N-term were considered as variable modifications. False discovery rate (FDR) was assessed using Percolator and thresholds for protein, peptide and modification site were specified at 1%. Label-free quantification (AUC) was performed within Proteomes Discoverer 2.5 and all protein abundances were normalized the CytC area.

### Native gel analysis

2 μg of protein samples (Strep-Tsa1 homooligomer, His_6_-Tsa2-EPEA homooligomer and Strep-Tsa1−His_6_-Tsa2-EPEA heterooligomer) were loaded in a 4-16% precast native gel (Serva), and run at 50 V for 10 min, followed by 180 V for 3 h. The protein ladder used was Native Marker Liquid Mix for BN/CN PAGE (Serva). Gel was stained with Coomassie InstantBlue® Protein Stain (Abcam) for 1 h at 4°C, destained with 20% (v/v) ethanol, 5% (v/v) acetic acid, and 1% (w/v) glycerol for 3 h at 4°C. Gel was stored in 5% glycerol at 4°C.

### Construction of yeast strains

All yeast strains used in this study are listed in **Supplementary Table 3**. The *TSA1*::*ROGFP2-TSA1*, *TSA1*::*ROGFP2-TSA1* Δ*tsa2*, *TSA2*::*ROGFP2-TSA2* and *TSA2*::*ROGFP2-TSA2* Δ*tsa1* strains were generated by standard homologous recombination approaches ^64^. *TSA1* and *TSA2* genes were first replaced by a URA3 cassette with a selection on Hartwell’s Complete (HC) agar plates lacking uracil. Subsequently the URA3 cassette was replaced by *ROGFP2-TSA1* or *ROGFP2-TSA2* with selection on HC plates containing 0.1% w/v 5-fluoroorotic acid (5-FOA, Zymo Research). 5-FOA is converted into toxic 5-flurouracil in cells harboring a functional URA3 and is therefore used as a negative-selection. Plates were incubated at 30°C for 48 hours. Colonies were picked and screened by PCR.

### Yeast peroxiredoxin hetero-oligomer induction experiments

Yeast strains were grown at 30°C as pre-cultures in HC Complete medium for 24 hours. Pre-cultures were diluted to an *D*_600_ = 1 in fresh HC Complete medium and grown for one hour. Cultures were then treated for the indicate timepoints with 1 mM H_2_O_2_. At the indicated timepoints, samples were for Native-PAGE analysis. For the Native-PAGE analysis, 25 *D*_600_ units were harvested by centrifugation at 800x*g* for 3 mins at room temperature. Cells were resuspended in 350 µL lysis buffer (50 mM Tris, pH7.7, 50 mM NaCl, 10 % (v/v) Glycerol, 20 mM NEM, 100 µM DTPA, 1 x protease inhibitor cocktail) and lysed by glass-bead homogenization. Lysates were cleared by centrifugation at 15,000x*g* for 15 mins at 5°C and the supernatant collected. Protein concentration in the supernatant was determined by Bradford assays and 20 µg protein was loaded per well of a 3–12% Clear-Native gel. After running gels were imaged for GFP fluorescence. Cell lysates were also analysed for total GFP fluorescence using a BMG Labtech CLARIOstar plate-reader.

### RoGFP2-based hetero-oligomer activity assays in yeast

BY4742 Δ*tsa1*Δ*tsa2* cells were transformed with p415TEF and p416TEF plasmids for the expression of roGFP2-peroxiredoxin fusion constructs and unfused peroxiredoxins variants respectively (**Supplementary Table 2**). Cells were grown to late-logarithmic phase (*D*_600_ = 3–4) in HC medium lacking the appropriate amino acids for plasmid selection. Cells were harvested by centrifugation at 800x*g* for 3 min at room temperature and resuspended to a final concentration of 7.5 *D*_600_ units/mL. Cells were transferred to a flat-bottomed 96-well imaging plate (BD Falcon 353219), with 200 µL cell suspension per well. Fully oxidized and fully reduced controls were established by the addition of 20 mM diamide and 100 mM DTT respectively, which allow for determination of the degree of roGFP2 oxidation (OxD) ^65^. A BMG Labtech CLARIOstar plate-reader was used to monitor fluorescence at the roGFP2 excitation, 400±7.5 nm and 488±7.5 nm, and emission, 510±10 nm. The experiment was inititated by the addition of exogenous H_2_O_2_ at the indicated concentration. OxD roGFP2 was calculated according to Equation 1.

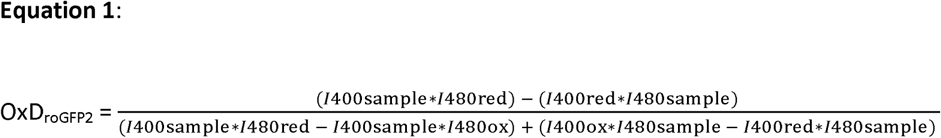

### Determination of the heterooligomeric state of mammalian PRDX1 and PRDX2 by ion exchange chromatography

HEK293 FLP-In/T-REx WT, PRDX1 KO and PRDX2 KO cells were seeded on 10 cm cell dish and analysed when cells were confluent. For harvesting, dishes were transferred to ice, washed twice with 10 mL of ice-cold AEC binding buffer (20 mM Tris, 20 mM NaCl, pH = 9.2) and harvested by scraping the cells into 2 mL of AEC binding buffer. Cells were lysed by sonication, cell debris was pelleted (20,000g for 30 minutes at 4°C), and the supernatant was loaded on a HiTrap Q HP anion exchange column (Cytiva) using an ÄKTA Start chromatography system. The column was previously equilibrated with 20 column volumes (CV) of AEC binding buffer and the bound lysate was washed with 20 CV of AEC binding buffer. Bound proteins were eluted from the column using a gradient (0-100% within 15 ml) of AEC elution buffer (20 mM Tris, 250 mM NaCl, pH = 9.2). A total of 30 eluted fractions with a volume of 0.5 ml were collected. All fractions were precipitated using 10% TCA (v/v) and stored at-80°C overnight. The next day, samples were thawed, precipitated proteins were pelleted (20,000g for 30 minutes at 4°C) and washed twice with ice-cold acetone. The acetone was completely removed by evaporation and the protein pellets were solubilized in 75 µl reducing LÄMMLI buffer. All samples were heat denatured and analyzed by SDS-PAGE and Western Blot. For final analysis, every second fraction was loaded.

### Bioinformatics analysis

Peroxiredoxin candidates were identified using iterative BLASTp (NCBI v2.16.0) with BLOSUM45 on the non-redundant protein database (GenPept, Swiss-Prot, PIR, PDF, PDB, RefSeq; Jan 9, 2025), restricted to genomes at assembly level “chromosome” or “complete.” New queries were generated from top-scoring hits in phylogenetic adjacent eukaryotic taxa to broaden and ensure coverage. Sequences containing characteristic peroxiredoxin PFAM domains (PF00578, PF08534, PF10417), identified via HMMER, were retained, and highly similar entries (>99% identity) within the same species were collapsed. Subcellular localization was predicted using DeepLoc 2.1 with default parameters.

## Acknowledgements

B.M. and M.D. gratefully acknowledge funding from the Deutsche Forschungsgemeinschaft (DFG, German Research Foundation) through the grants MO 2774/6-1 project number 505680640, MO 2774/7-1 and DE 1431/19-1 project number 508372800. J.M. was supported by a VIB grant. J.M.P. was supported by a FWO fellowship (1193524N). The DFG funds research in the Laboratory of JR through the grants RI2150/5-1 project number 435235019, RTG2550/2 project number 411422114, SPP2453 project number 541742459, CRC1218 - project number 269925409, and CRC1678 – project number 520471345. A.S. acknowledges funding from the COMPETE 2020 - Operational Programme for Competitiveness and Internationalisation and Portuguese national funds via FCT – Fundação para a Ciência e a Tecnologia, under projects UIDB/04539/2020, UIDP/ 04539/2020, LA/P/0058/2020, UIDB/00313/2020, and UIDP/00313/ 2020. We thank Dr. Els Pardon (Steyaert Lab, VIB-VUB Center for Structural biology) for kindly providing the Nbsyn2.20 nanobody. We thank Juhans Dechenne (Louvain Drug Research Institute, The Medicinal Chemistry Group, UCL) for his help with nanoDSF experiments. We thank the BECM VIB-VUB cryo-EM imaging facility in Brussels and Dr. Marcus Fislage for the support during negative staining EM imaging and processing. AMT acknowledges support from National Funds through FCT—Fundação para a Ciência e a Tecnologia, I.P., under the project UIDB/04293/2020.

