## Supplementary Information for "Heterooligomerization drives structural plasticity of eukaryotic peroxiredoxins"

### for

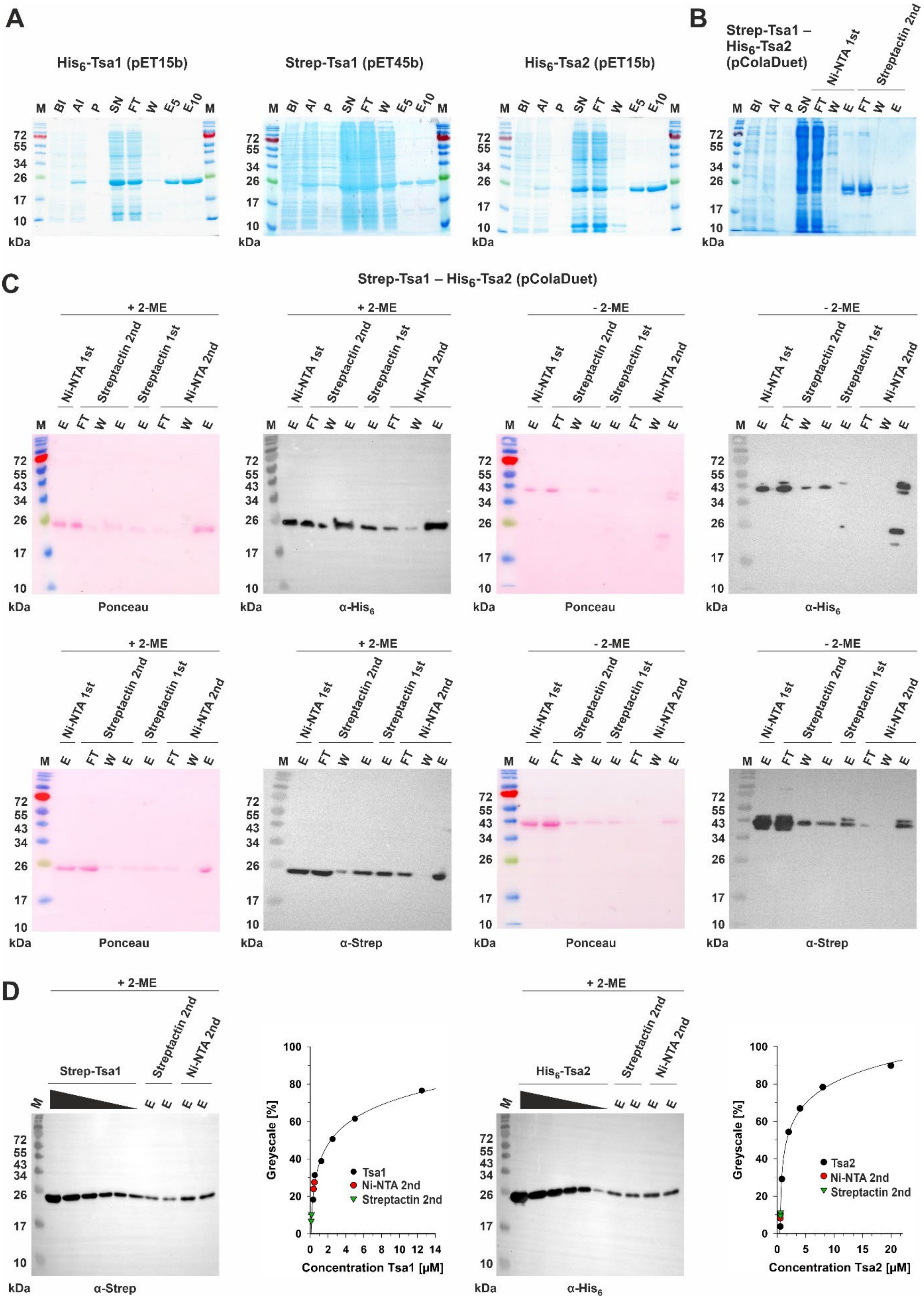
A
His$_6$-Tsa1 (pET15b)
Strep-Tsa1 (pET45b)
His$_6$-Tsa2 (pET15b)
M BI AI P SN FT W E5 E10 M
72 55 34 26 17 10
kDa
B
Strep-Tsa1 – His$_6$-Tsa2 (pColaDuet)
Ni-NTA 1st
Streptactin 2nd
M BI AI P SN FT W E FT W E
C
Strep-Tsa1 – His$_6$-Tsa2 (pColaDuet)
+ 2-ME
- 2-ME
Ni-NTA 1st
Streptactin 2nd
Streptactin 1st
Ni-NTA 2nd
M E FT W E E FT W E
72 55 43 34 26 17 10
kDa
Ponceau
α-His$_6$
α-Strep
D
+ 2-ME
Strep-Tsa1
His$_6$-Tsa2
Streptactin 2nd
Ni-NTA 2nd
M E E E E
α-Strep
α-His$_6$
Greyscale [%]
Concentration Tsa1 [µM]
Concentration Tsa2 [µM]
Tsa1
Tsa2
Ni-NTA 2nd
Streptactin 2nd

**Supplementary Figure 1. Purification of recombinant Tsa1 and Tsa2 from *E. coli*.**

**A**) SDS-PAGE analysis of representative individual purifications of recombinant $His_6$-Tsa1 (left), Strep-Tsa1 (middle), and $His_6$-Tsa2. **B**) SDS-PAGE analysis of a representative tandem affinity co-purification of recombinant Strep-Tsa1 and $His_6$-Tsa2. **C**) Western blot analysis of the tandem affinity co-purifications of recombinant Strep-Tsa1 and $His_6$-Tsa2. Samples were separated by reducing (left) or non-reducing (right) SDS-PAGE and blots were stained with ponceau as a loading control. The membranes were subsequently decorated with an antibody against either the His-tag (upper row) or the Strep-tag (lower row). Each membrane shows two tandem affinity co-purifications, one with Ni-NTA agarose followed by StrepTactin agarose and one with StrepTactin agarose followed by Ni-NTA agarose. **D**) Semi-quantitative western blot analysis of the protein content of co-purified Strep-Tsa1 and $His_6$-Tsa2. Known concentrations of individually purified Strep-Tsa1 or $His_6$-Tsa2 were used for calibration. Greyscales were quantified using ImageJ. The calculated molecular masses of $His_6$-Tsa1/2 and Strep-Tsa1 are 23.8 and 22.9 kDa, respectively. M, marker; BI, before induction; AI, after induction; P, pellet after sonication; SN, supernatant; FT, flow-through; W, wash; $E_{(5,10)}$, eluate (5 or 10 µL loaded); 2-ME, 2-mercaptoethanol.

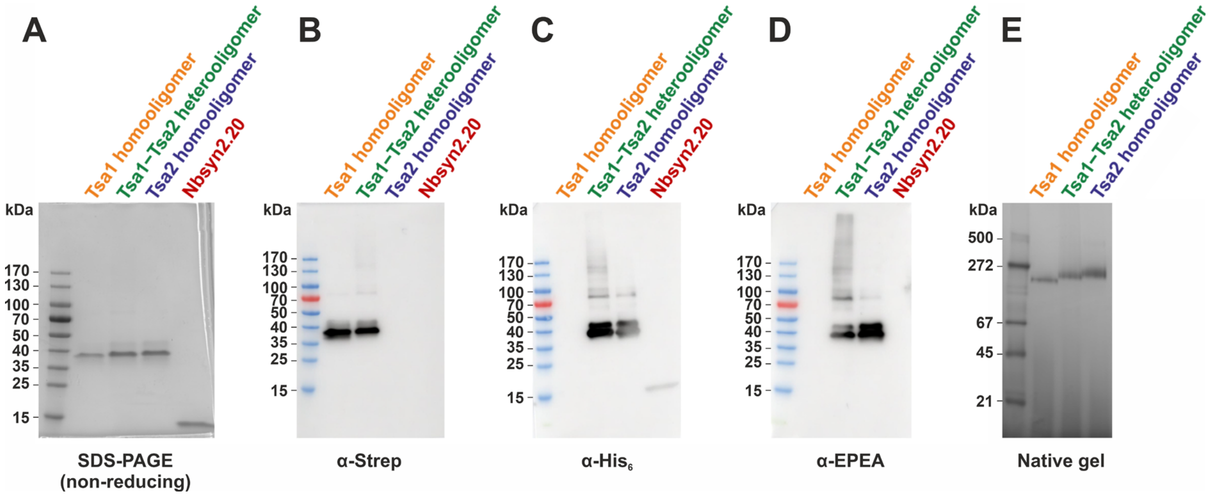


**Supplementary Figure 2. Purified proteins present the correct tags identified by western blot**

Purified Strep-Tsa1 homooligomer, $His_6$-Tsa2-EPEA homooligomer, 0.88 mol/mol Strep-Tsa1–$His_6$-Tsa2-EPEA heterooligomer and His-Nbsyn2.20 samples were separated on a 4–20% gradient SDS-PAGE gel and transferred onto PVDF membranes. The membranes were probed with anti-EPEA, HRP-conjugated anti-Strep and anti-His primary antibodies. Then, anti-EPEA and anti-His membranes were incubated with HRP-conjugated goat anti-mouse IgG secondary antibody. Protein bands were visualized with HRP substrate Pierce™ ECL western blotting substrate. **A)** 4-20% SDS-PAGE gradient gel **B)** anti-Strep blot **C)** anti-$His_6$ blot **D)** anti-EPEA blot **E)** Native gel analysis shows that Strep-Tsa1–$His_6$-Tsa2-EPEA heterooligomers form a single stable population in solution. 2 µg of protein samples (Strep-Tsa1 homooligomer, $His_6$-Tsa2-EPEA homooligomer, 0.88 mol/mol Strep-Tsa1–$His_6$-Tsa2-EPEA heterooligomer) were separated on a 4-16% precasted native gel (Serva). The protein ladder used was Native Marker Liquid Mix for BN/CN PAGE (Serva).

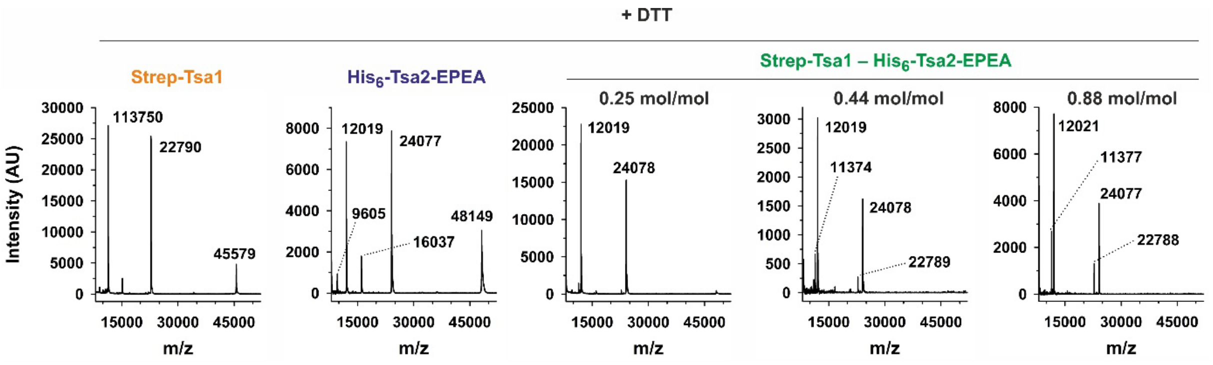


**Supplementary Figure 3. Purified Strep-Tsa1 homooligomer, His$_6$-Tsa2-EPEA homooligomer, 0.88 mol/mol Strep-Tsa1–His$_6$-Tsa2-EPEA heterooligomer samples do not show any contamination with AhpC from *E. coli***

MALDI-TOF mass spectrometry analysis in reducing conditions reveals that Strep-Tsa1–His$_6$-Tsa2-EPEA heterooligomers are formed by Strep-Tsa1 and His$_6$-Tsa2-EPEA monomers in varying ratios. The ratio of each Strep-Tsa1–His$_6$-Tsa2-EPEA heterooligomer sample is displayed above the spectra. The intensity of the signal (AU) is shown at different m/z ratios. Duplicate measurements were performed for each sample, and data were analyzed by FlexAnalysis 3.4 (Bruker).

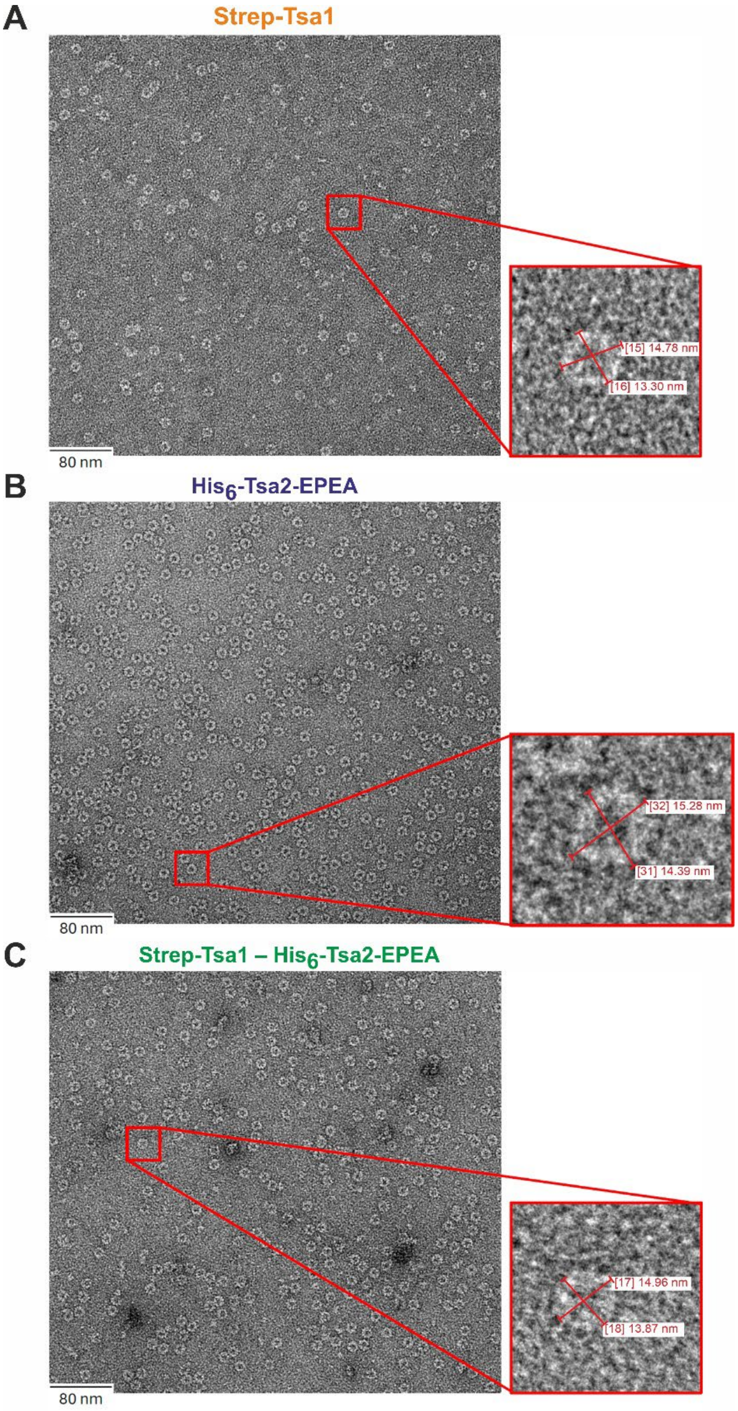


**Supplementary Figure 4. The presence of the N- and C-terminal tags does not affect the decameric stability of the proteins in solution**

Protein particles appear bright against the 2% uranyl-acetate stain, display a characteristic decameric 'donut-like' structure. Diameters were measured using EMMenu from TVIPS imaging software. Protein concentration was 0.02 mg/mL. **A)** Strep-Tsa1 homooligomer **B)** $His_6$-Tsa2-EPEA homooligomer **C)** Strep-Tsa1–$His_6$-Tsa2-EPEA heterooligomer

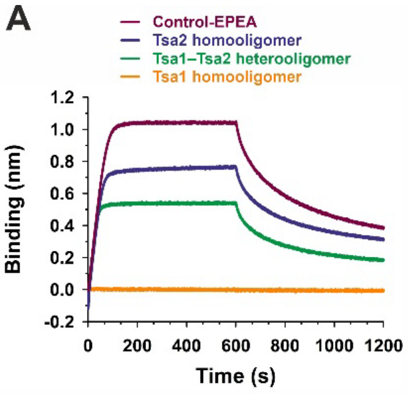


| | $K_D$ (nM) | $k_a$ ($\mu M^{-1}s^{-1}$) | $k_d$ ($s^{-1}$) |
|---|---|---|---|
| Control-EPEA | 49.4 ± 9.5 | 0.2 ± 0.2 | 0.01 ± $10^{-4}$ |
| Tsa2 homooligomer | 49.0 ± 17.7 | 0.3 ± 0.1 | 0.01 ± $10^{-4}$ |
| Tsa1–Tsa2 heterooligomer | 23.9 ± 4.9 | 0.5 ± 0.2 | 0.01 ± $10^{-4}$ |
| Tsa1 homooligomer | - | - | - |

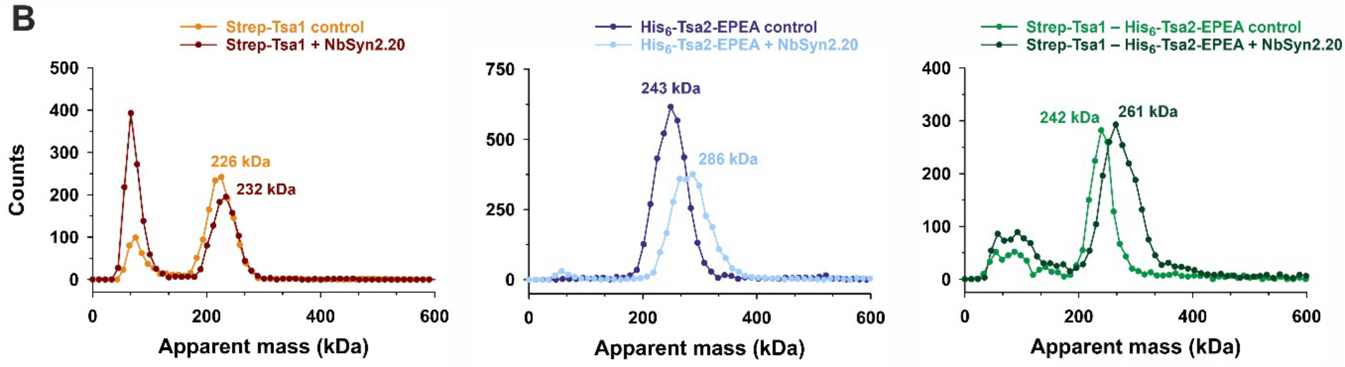


| | Apparent Mass (kDa) ± SD |
|---|---|
| Tsa1 homodecamer control | 226 ± 2.5 |
| Tsa1 homodecamer + NbSyn2.20 | 232 ± 3.1 |
| Tsa2 homodecamer + control | 243 ± 6.6 |
| Tsa2 homodecamer + NbSyn2.20 | 286 ± 6.8 |
| Tsa1 – Tsa2 heterodecamer control | 242 ± 4.0 |
| Tsa1 – Tsa2 heterodecamer NbSyn2.20 | 261 ± 4.7 |

**Supplementary Fig. 5. Nbsyn2.20 specifically binds to the C-terminal EPEA-tag of $His_6$-Tsa2 protein**

**A)** BLI assay shows that Nbsyn2.20 specifically binds to the C-terminal EPEA-tag of Tsa2. Strep-Tsa1 homooligomer, $His_6$-Tsa2-EPEA homooligomer, and 0.88 mol/mol Strep-Tsa1–$His_6$-Tsa2-EPEA heterooligomer were immobilized on SA-sensors, with His-Nbsyn2.20 used as the analyte. Protein-EPEA was used as a positive control. Association and dissociation curves are shown (Binding (nm) vs time (s)) with Strep-Tsa1 homooligomer, $His_6$-Tsa2-EPEA homooligomer, 0.88 mol/mol Strep-Tsa1–$His_6$-Tsa2-EPEA heterooligomer and protein-EPEA represented in orange, blue, green and purple, respectively. Data were analyzed using Octet Analysis studio 13.0 software, and the graph was generated with GraphPad Software, Inc. Triplicates were measured. **B)** Mass photometry experimental data show that at least two Nbsyn2.20 molecules bind to the 0.88 mol/mol Strep-Tsa1–$His_6$-Tsa2-EPEA heterooligomer. Stock protein solutions (5 mM) were diluted prior to the measurement. Data were acquired for 60 s, and counts of individual molecules were plotted against their molecular weight. Strep-Tsa1 and $His_6$-Tsa2-EPEA homooligomers mass photometry histograms are shown in orange and blue, respectively, while the histogram for 0.88 mol/mol Strep-Tsa1–$His_6$-Tsa2-EPEA heterooligomer is shown in green. Data were analyzed using DiscoverMP (version 2.1.1; Refeyn Ltd). Triplicates were measured, and the average molecular weight was calculated.

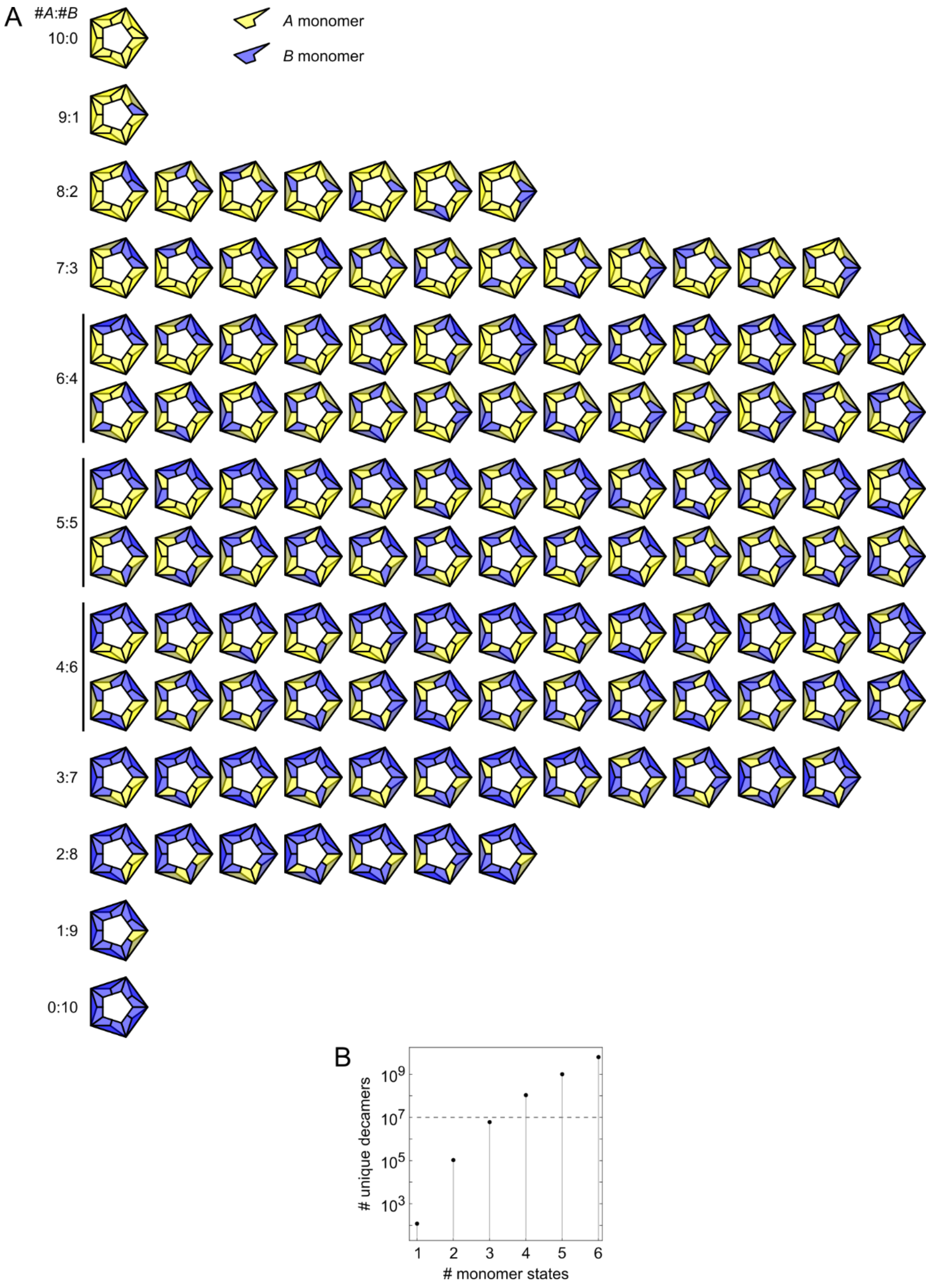


**Supplementary Figure 6. Enumeration of the distinct decamers that can be formed from two types of monomers**

**A**) Distinct decamers as a function of the stoichiometric ratio between monomer types. **B**) Number of unique hetero-decamer configurations as a function of the number of possible states each monomer type can adopt. The horizontal dashed line indicates the estimated number of peroxiredoxin decamers in a typical mammalian cell. Note the logarithmic scale on the vertical axis. Calculations are explained here below.

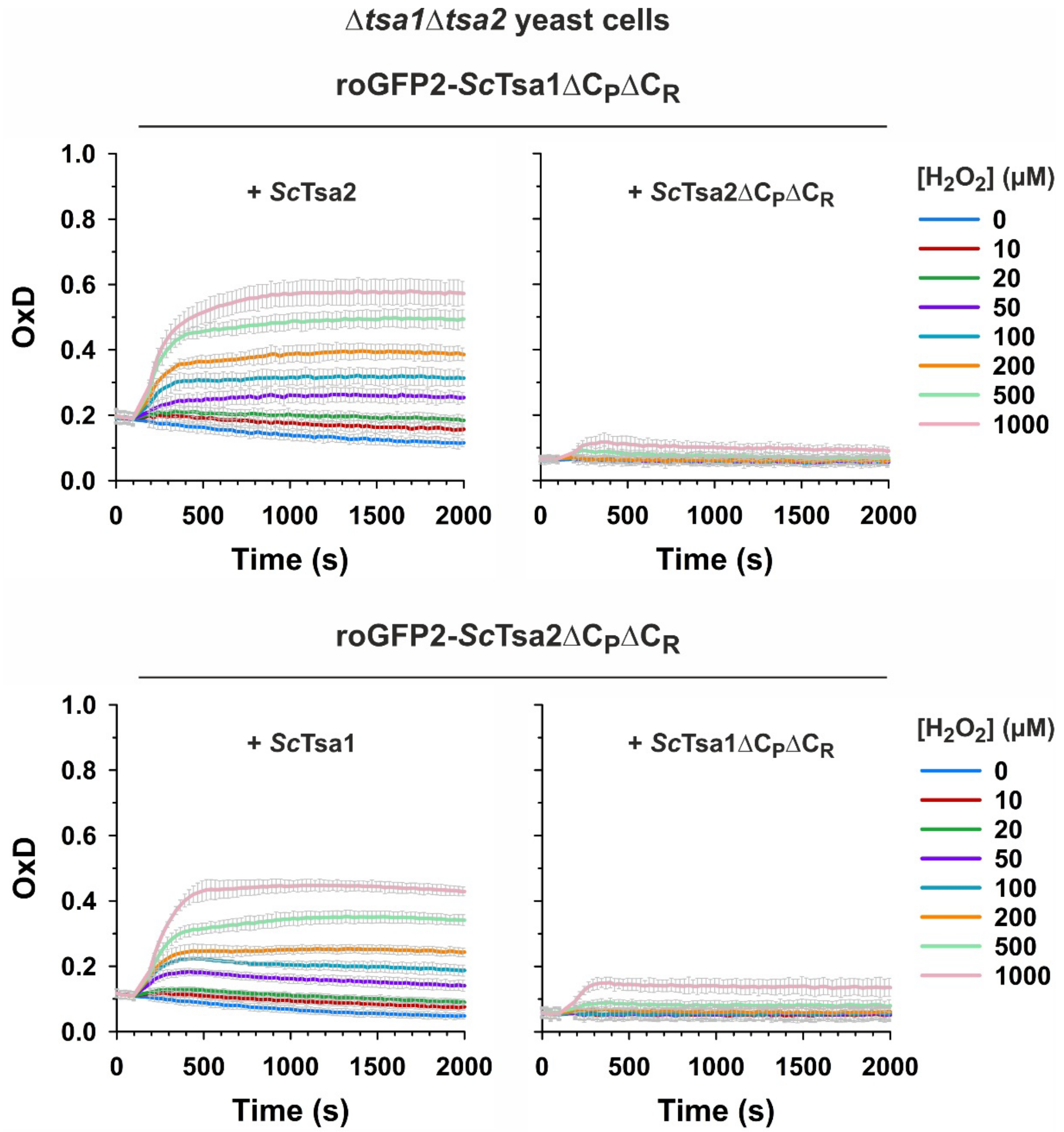


**Supplementary Figure 7. Tsa1 and Tsa2 form enzymatically active hetero-oligomers in yeast**

Graphs showing the change in the degree of oxidation (OxD) in response to 1 mM $H_2O_2$ of roGFP2-Tsa1Δ$C_P$Δ$C_R$ and roGFP2-Tsa1Δ$C_P$Δ$C_R$ constructs expressed in Δ*tsa1*Δ*tsa2* yeast together with either a wild-type (wt) or cysteine-less (Δ$C_P$Δ$C_R$) variant of the corresponding partner peroxiredoxin. Experiments were repeated 3 times with independent yeast cultures. Data are presented as mean ± s.d.

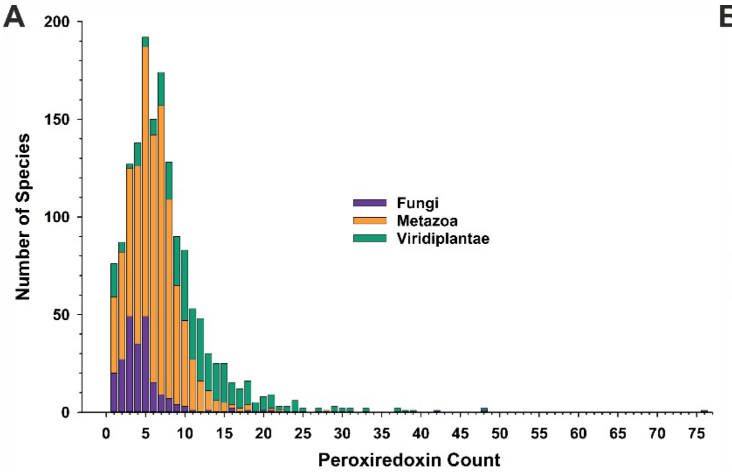


**Supplementary Figure 8. Multiple Prx1/AhpC-type peroxiredoxin isoforms are found within at least one subcellular compartment in most eukaryotes.**

Corresponds to main Figure 7C. A) Histogram showing the total number of Prx1/AhpC-type peroxiredoxins predicted within each of 1525 sampled eukaryotic species. **B**) Histogram showing the number of species in which there are predicted to be the indicated number of Prx1/AhpC-type peroxiredoxin present within the indicated subcellular compartments.

**Supplementary Table 1. Primers used in this study**

| **Plasmid** | **Encoded Protein** | **Primers for cloning/mutagenesis** |
|---|---|---|
| pET15b/His-*TSA1* | MGSS$H_6$SSGLVPRGSHM-Tsa1 | **Ref. [27]** |
| pET15b/His-*TSA2* | MGSS$H_6$SSGLVPRGSHM-Tsa2 | **Ref. [27]** |
| pET45b/Strep2-*TSA1* | MAWSHPQFEKGGT-Tsa1 | S: GATC**GGTACC**GTCGCTCAAGTTCAAAAGCAAG<br>AS: GATC**CCTAGG**TTATTTGTTGGCAGCTTCGAAG |
| pColaDuet/Strep2-*TSA1*/His-*TSA2* | MAWSHPQFEKGGT-Tsa1<br>MGSS$H_6$SSGLVPRGSHM-Tsa2 | S: GATC**CATATG**GCATGGTCTCATCCGCAGTTTG<br>AS: GATC**CTCGAG**TTATTTGTTGGCAGCTTCGAAG |
| pET45b/Strep2-*TSA1ΔC$_R$* | MAWSHPQFEKGGT-Tsa1*ΔC$_R$* | S: GGTACTGTCTTGCCATCTAACTGGACTCCAGGTG<br>AS: CACCTGGAGTCCAGTTAGATGGCAAGACAGTACC |
| pColaDuet/Strep2-*TSA1ΔC$_R$* /His-*TSA2* | MAWSHPQFEKGGT-Tsa1*ΔC$_R$*<br>MGSS$H_6$SSGLVPRGSHM-Tsa2 | - |
| pColaDuet/Strep2-*TSA1*/His-*TSA2-EPEA* | MAWSHPQFEKGGT-Tsa1<br>MGSS$H_6$SSGLVPRGSHM-Tsa2-EPEA | S: GATC**CCATGG**GCAGCAGCCATCATCATCATCATCACAGCA<br>AS: GATC**GGATCC**TTATGCTTCCGGTTCATTATTGGCATTTTTGAAATACTCC |
| pET15b/His-*TSA2-EPEA* | MGSS$H_6$SSGLVPRGSHM-Tsa2-EPEA | **-** |
| pTrc99aNHis-ScTRX1 | M$H_6$P-ScTrx1 | **Ref. [27]** |

**Supplementary Table 2. Yeast strains used in this study.**

| Genotype | Source |
|---|---|
| BY4742 *MATα his3Δ1 leu2Δ1 lys2Δ0 ura3Δ0* | Euroscarf |
| BY4742 *TSA1*::*ROGFP2-TSA1* | This study |
| BY4742 *TSA1*::*ROGFP2-TSA1* Δ*tsa2*::*kanMX4* | This study |
| BY4742 *TSA2*::*ROGFP2-TSA2* | This study |
| BY4742 *TSA2*::*ROGFP2-TSA1* Δ*tsa1*::*kanMX4* | This study |
| BY4742wt +p416TEF roGFP2-Tsa1Δ$C_P$Δ$C_R$ + p415TEF Tsa2 | This study |
| BY4742wt +p416TEF roGFP2-Tsa1Δ$C_P$Δ$C_R$ + p415TEF Tsa2Δ$C_P$Δ$C_R$ | This study |
| BY4742wt +p416TEF roGFP2-Tsa2Δ$C_P$Δ$C_R$ + p415TEF Tsa1 | This study |
| BY4742wt +p416TEF roGFP2-Tsa2Δ$C_P$Δ$C_R$ + p415TEF Tsa1Δ$C_P$Δ$C_R$ | This study |
| BY4742wt +p416TEF roGFP2-*Hs*PRDX1Δ$C_P$Δ$C_R$ + p415TEF *Hs*PRDX2 | This study |
| BY4742wt +p416TEF roGFP2-*Hs*PRDX1Δ$C_P$Δ$C_R$ + p415TEF *Hs*PRDX2Δ$C_P$Δ$C_R$ | This study |
| BY4742wt +p416TEF roGFP2-*Hs*PRDX2Δ$C_P$Δ$C_R$ + p415TEF *Hs*PRDX1 | This study |
| BY4742wt +p416TEF roGFP2-*Hs*PRDX2Δ$C_P$Δ$C_R$ + p415TEF *Hs*PRDX1Δ$C_P$Δ$C_R$ | This study |
| BY4742wt +p416TEF roGFP2-BAS1AΔ$C_P$Δ$C_R$ + p415TEF BAS1B | This study |
| BY4742wt +p416TEF roGFP2-BAS1AΔ$C_P$Δ$C_R$ + p415TEF BAS1BΔ$C_P$Δ$C_R$ | This study |
| BY4742wt +p416TEF roGFP2-BAS1BΔ$C_P$Δ$C_R$ + p415TEF BAS1A | This study |
| BY4742wt +p416TEF roGFP2-BAS1BΔ$C_P$Δ$C_R$ + p415TEF BAS1AΔ$C_P$Δ$C_R$ | This study |
| BY4742wt +p416TEF roGFP2-*Li*PRDX1Δ$C_P$Δ$C_R$ + p415TEF *Li*PRDX2 | This study |
| BY4742wt +p416TEF roGFP2-*Li*PRDX1Δ$C_P$Δ$C_R$ + p415TEF *Li*PRDX2Δ$C_P$Δ$C_R$ | This study |
| BY4742wt +p416TEF roGFP2-*Li*PRDX2Δ$C_P$Δ$C_R$ + p415TEF *Li*PRDX1 | This study |
| BY4742wt +p416TEF roGFP2-*Li*PRDX2Δ$C_P$Δ$C_R$ + p415TEF *Li*PRDX1Δ$C_P$Δ$C_R$ | This study |
| BY4742wt +p416TEF roGFP2-BAS1A + p415TEF Empty | This study |
| BY4742wt +p416TEF roGFP2-BAS1A + p415TEF BAS1A | This study |
| BY4742wt +p416TEF roGFP2-BAS1A + p415TEF BAS1B | This study |
| BY4742wt +p416TEF roGFP2-BAS1B + p415TEF Empty | This study |

| | |
|---|---|
| BY4742wt +p416TEF roGFP2-BAS1B + p415TEF BAS1A | This study |
| BY4742wt +p416TEF roGFP2-BAS1B + p415TEF BAS1B | This study |
| BY4742wt +p416TEF roGFP2-*Li*PRDX1 + p415TEF Empty | This study |
| BY4742wt +p416TEF roGFP2-*Li*PRDX1 + p415TEF *Li*PRDX1 | This study |
| BY4742wt +p416TEF roGFP2-*Li*PRDX1 + p415TEF *Li*PRDX2 | This study |
| BY4742wt +p416TEF roGFP2-*Li*PRDX2 + p415TEF Empty | This study |
| BY4742wt +p416TEF roGFP2-*Li*PRDX2 + p415TEF *Li*PRDX1 | This study |
| BY4742wt +p416TEF roGFP2-*Li*PRDX2 + p415TEF *Li*PRDX2 | This study |

**Computing the number of unique decamer configurations**

*Unique decamers formed from two types of monomers, containing a given number of monomers of one type*

If each position in a peroxiredoxin decamer can be occupied by one of two distinct monomer types (*A*, *B*), then there are $2^{10}$ = 1024 possible configurations, of which $\binom{10}{a}$ have exactly $a$ monomers of type *A*. However, most of these configurations are equivalent under rotation around the 5-fold rotational symmetry axis perpendicular to the equatorial plane of the toroidal decamer (Fig. 1a), or the five 2-fold rotational symmetry axes in this plane and crossing the center of each dimer. To accurately enumerate the truly distinct decamer configurations, we apply the Cauchy–Frobenius lemma (a.k.a. Burnside's lemma [1]. This lemma asserts that the number of distinct configurations $(N)$ of a symmetric object is given by the formula:

$$N = \frac{1}{|G|}\sum_{g\in G} Fix(g), \tag{1}$$

where $G$ is the symmetry group of the object, which here includes rotations around the above-mentioned 5-fold and 2-fold axes; $|G|$ is the order of the group; and $Fix(g)$ is the number of configurations that are invariant under a given group element $g$.

In the present case, $|G| = 10,$ of which one element is the identity operation, four are the 72°, 144°, 216° and 288° rotations around the 5-fold axis, and the remaining five are the 180° rotations around the five 2-fold axes.

We must now calculate $Fix(g)$ for each of these symmetry operations. Because every configuration is invariant under the identity operation $(e)$, there are

$$Fix(e,a) = \binom{10}{a} \tag{2}$$

invariant configurations with exactly $a$ *A*-type monomers.

In turn, a configuration is only invariant under rotation around the 5-fold axis if all five dimers are identical. This is only possible for $a = 0$ (1 configuration: all dimers *BB*), $a = 5$ (2 configurations: all *AB* or all *BA*), and $a = 10$ (1 configuration: all *AA*). Therefore:

$$Fix(g_{\text{5-fold}}, a) = 4 \times \begin{cases} 1 & a \in \{0,10\} \\ 2 & a = 5 \\ 0 & \text{otherwise} \end{cases}, \tag{3}$$

where the factor 4 accounts for the possible rotations around this symmetry axis.

Turning attention to rotations around each 2-fold axis, configurations invariant under this operation must meet both the following conditions: (i) dimers at which the axis is centered must be either *AA* or *BB*, and (ii) each monomer in dimers to one side of the axis must match another one of the same type in the opposite side of the axis. This can only happen for even $a$, and since there are five independent *pairs* of monomers the number of such configurations is:

$$Fix(g_{2\text{-fold}}, a) = 5 \times \begin{cases} \begin{pmatrix} 5 \\ a/2 \end{pmatrix} & a\%2 = 0 \\ 0 & a\%2 = 1 \end{cases}, \qquad (4)$$

where the 5 factor accounts for the number of these axes.

Finally, the total number of configurations with exactly $a$ *A*-type monomers is obtained by replacing equations (2)–(4), according to equation (1):

$$N(a) = \frac{1}{10}\left(Fix(e,a) + Fix(g_{5\text{-fold}}, a) + Fix(g_{2\text{-fold}}, a)\right). \qquad (5)$$

This yields the values described in the main text, with representations of the unique configurations in **Supplementary Fig. 6**.

*Unique decamers formed of two types of monomers, each having n possible states*

If there are two types of monomers and each can adopt $n$ states, there is a total of $(2n)^{10}$ possible decamer configurations. In order to find how many of these are unique, we resort again to the Cauchy–Frobenius lemma, equation (1.1), without the constraints on the number of monomers of a given type. In this case, all the $(2n)^{10} = 1024n^{10}$ configurations are invariant under the identity operation. For invariance under the 5-fold rotations all dimers must be equal, which $4 \times (2n)^2 = 16n^2$ configurations satisfy. In turn, for invariance under the 2-fold rotations, both monomers in one of the dimers must be of the same type and state ($2n$ choices), and each of the remaining two pairs of dimers must be identical ($(2n)^2$ choices per pair). Therefore, for the five 2-fold rotations a total of $5 \times (2n) \times (2n)^2 \times (2n)^2 = 160n^5$ configurations are invariant. The number of unique configurations is thus:

$$N(n) = \frac{1024n^{10} + 160n^5 + 16n^2}{10}, \qquad (6)$$

which is plotted in **Supplementary Fig. 6b**.

### Supplementary References


1. Burnside, W. *Theory of Groups of Finite Order*, (Cambridge University Press, 1897).
